# A molecular map of the living human brain from quantitative MRI

**DOI:** 10.1101/2025.09.24.678316

**Authors:** Melissa Grant-Peters, George E C Thomas, Daan van Kruining, Louise A Huuki-Myers, Ruth Zhang, Jonathan W Brenton, Hemanth Nelvagal, Angelika Zarkali, James R Evans, Christina E Toomey, Aine Fairbrother-Browne, Ivelina Dobreva, Kelsey D Montgomery, Joanne Lachica, Naomi Hannaway, Nicholas Wood, Sonia Gandhi, Martina F Callaghan, Frances Platt, Zane Jaunmuktane, Karin Shmueli, Leonardo Collado-Torres, Kristen R Maynard, Rimona S Weil, Mina Ryten

**Author notes:** These authors contributed equally (Melissa Grant-Peters, George E C Thomas, Rimona S Weil, Mina Ryten).

## Abstract

Profiling dynamic molecular processes in neurodegeneration in vivo remains a major clinical challenge. Limited brain tissue accessibility especially limits the development and effective deployment of emerging disease modifying therapies, highlighting the need for non-invasive methods for profiling molecular disease processes. We introduce a non-invasive imaging framework that integrates ultra-high-resolution 7T quantitative MRI (qMRI) with spatially-resolved transcriptomics (SRT) to infer cell-type and pathway-specific molecular features within the cortical grey matter. This proof-of-concept establishes the integration of qMRI and SRT as a scalable, non-invasive platform for molecular profiling in neurodegeneration, with significant potential for precision therapeutic monitoring, drug development and clinical trials.

## Introduction

The advent of high-resolution and high-throughput molecular profiling techniques is providing unprecedented insight into the processes underlying neurodegeneration. Consequently, the field is seeing growth in the development of drugs targeting specific molecular features, which are considerably more likely to succeed in clinical trials than non-targeted therapies. This is showcased by the emergence of disease modifying treatments (DMTs) for Alzheimer’s disease (*1*) and amyotrophic lateral sclerosis (*2*). However, it is critical for successful deployment of these drugs that there is an effective and scalable strategy for patient stratification based on the molecular features of interest. Diseases affecting tissues which can easily be biopsied for molecular classification such as certain types of cancer (*3*, *4*) and autoimmune disorders (*5*) showcase the value of molecular classification for drug development, clinical trials as well as therapeutic decision-making. In contrast, in neurodegenerative disorders, the ethical concerns and inaccessibility of brain tissue significantly limit drug development, patient stratification in drug trials and streamlining of therapeutic decision making, highlighting the urgent need for precision medicine tools in neurodegeneration and neurology more generally (*6*, *7*).

Meeting this need requires a precision medicine framework that is both patient-centred and compatible with the existing clinical infrastructure. Such an approach must ensure tolerability (favouring non-invasive methods over biopsies or high-risk procedures), enable simultaneous profiling of multiple molecular features to reduce cost and procedural burden, and maximise the utility of resources already in place. Strategies that fail to meet these criteria can struggle to achieve clinical translation; for example, lecanemab for Alzheimer’s disease (*1*) requires early-stage confirmation via positron emission tomography (PET) imaging, but the limited availability of PET infrastructure has constrained its population-level impact.

Magnetic resonance imaging (MRI) offers a way forward, as it satisfies many of these requirements by providing a non-invasive, widely accessible, and clinically established platform. So far, MRI has proven indispensable in capturing structural features of disease. For example, in multiple sclerosis MRI is an essential tool in diagnosis, disease monitoring, and treatment decisions (*8*), illustrating how imaging can successfully anchor a precision medicine framework in neurology. MRI has also proven valuable for assessing neurodegenerative disorders such as Alzheimer’s disease, with structural MRI yielding spatially resolved insights into disease processes by tracking anatomical changes across affected brain regions (*9*). Similarly, it can be used to evaluate Parkinson’s disease progression, since disease progression follows a region-specific trajectory (*10*, *11*). However, such structural changes in neurodegeneration only emerge after profound and irreversible cellular damage, highlighting the need for methods which can detect earlier molecular features (**Fig 1A**).

**Figure 1.**
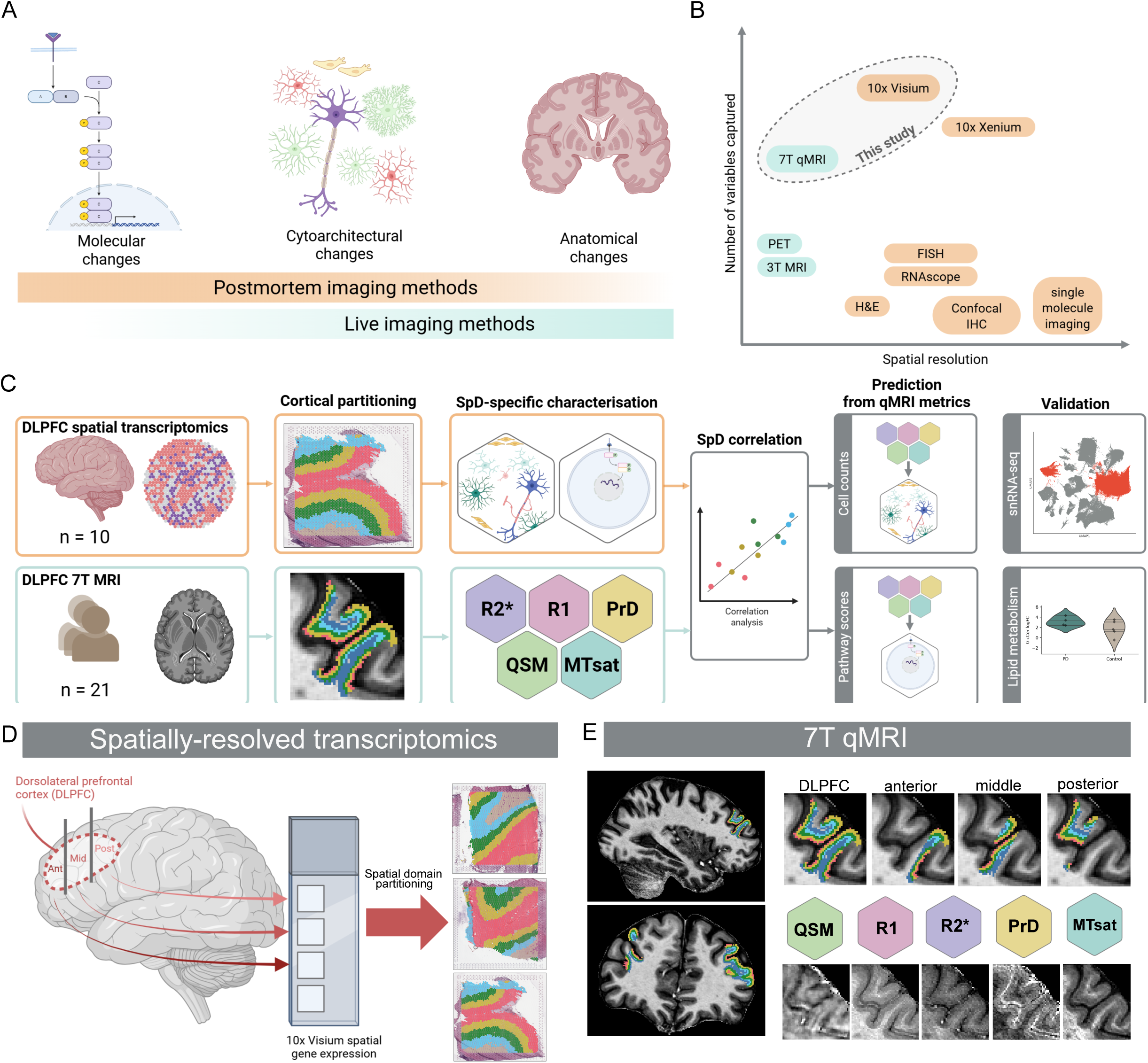
Integration of ultra-high resolution qMRI and SRT. (**A**) MRI technologies can capture non-invasively disease features in live patients, however by the time disease features are detectable anatomically, irreversible cellular and molecular damage has already occurred. In contrast, spatially-resolved transcriptomics (SRT) methods can capture molecular and cellular features which might be present in early disease, but are generally employed in postmortem samples. (**B**) Recent advances in both qMRI and SRT mean the resolution and number of variables captured by these variables make integration of these methods possible for the first time. (**C**) Diagram of sample structure of the DLPFC. Samples were dissected to include anterior, middle and posterior DLPFC. Grey matter of each section was partitioned into 4 spatial domains. (**D**) Diagram of sample structure of 7T qMRI data. DLPFC was selected according to region selection of SRT data and grey matter cortex was partitioned into 4 spatial domains. (**E**) Study design diagram. We used SRT data to calculate estimates of cellular composition and expression of biological pathways in each ROI. We characterized the relationship of these metrics in relation to qMRI metrics from these ROIs in 7T qMRI. We then created models to predict the molecular and cellular information from qMRI data.

Quantitative MRI (qMRI) has significant untapped potential to reveal subtle molecular changes which occur before widespread structural alterations are found (*12–14*). qMRI extends the utility of MRI capturing molecular-level features through advanced modelling of imaging data obtained using widely available scanner technology. In contrast to blood and cerebrospinal fluid (CSF) biomarkers, which also capture molecular-level features, qMRI is non-invasive and spatially-resolved. It is increasingly gaining traction, with techniques such as multi-parameter mapping (MPM) enabling the extraction of a wide range of continuous, quantitative variables across the whole brain (*15*). These provide insight into tissue microstructure and potentially capture dynamic changes across the disease course, from prodromal to late stages (*16–19*). In contrast with methods such as PET imaging, which allows for the targeting of specific molecular disease features, qMRI is more widely available (*20*), can capture multiple metrics, is non-invasive and does not entail any ionising radiation dose. Moreover, the deployment of novel MR imaging sequences is significantly faster than the deployment of novel PET radiotracers, which requires several additional steps including regulatory approval (*21*, *22*). Despite this, qMRI, including MPMs, remains relatively underutilized clinically and is typically employed as an adjunct to conventional structural imaging in research settings (*23*).

Increasing interpretability of qMRI has transformative potential. Specifically, qMRI captures signal from a complex mixture of cellular and molecular sources (*24*, *25*). Deconvoluting the contributions of these cellular and molecular entities in qMRI signals would facilitate leveraging of qMRI in clinical contexts (*26*, *27*), particularly as DMTs increasingly target specific cellular and molecular mechanisms. Spatially-resolved transcriptomics (SRT) are methods which hold significant potential for this purpose. These methods, which enable the mapping of gene expression data in up-to single-cell resolution, enable untargeted quantification of tens of thousands of RNA species from a single tissue section on a two dimensional plane. At the same time, recent improvements in ultra-high-field MRI (≥7T) deliver sufficient spatial resolution to provide scope for spatial integration of qMRI with SRT by aligning microstructural imaging with molecular data. Importantly, due to the spatially resolved nature of both datasets, microstructural and anatomical landmarks can be used to integrate information across separate cohorts. This convergence creates a timely opportunity to resolve the biological bases of qMRI metrics and unlock their potential for precision medicine in neurology (**Fig 1B**).

In this work, we show that qMRI can be molecularly interpreted using SRT, enabling non-invasive inference of laminar molecular pathology in living humans. We demonstrate that molecular pathways and cell type presence have significant correlative relationships with qMRI metrics, and that disruptions to these biological entities in disease can be predicted with qMRI data. This means that qMRI can be applied to living individuals to infer molecular and cell type changes that were previously only measurable with transcriptomics data from postmortem donors. Our approach has especially high potential for disease staging and patient stratification for clinical trials, and to provide mechanistic insights into disease heterogeneity and pathophysiology in living patients.

## Results

### Shared spatial framework of cortical spatial domains in qMRI and SRT

Our live imaging cohort consisted of 21 healthy controls (age = 67.7 ± 7.96 years, 10/21 females, **ST1**). MPM sequences were acquired on a 7T MRI system at 0.6 mm isotropic resolution. From these, we generated quantitative maps of magnetic susceptibility (QSM), proton density (PrD), longitudinal relaxation rate (R_1_), effective transverse relaxation rate (R_2_*), and magnetization transfer saturation (MTsat). Each of these metrics is sensitive to differing microstructural features in the tissue. In brief, in the brain, QSM is primarily sensitive to paramagnetic iron and diamagnetic myelin that have opposing influences (*28*, *29*); R_1_ to myelin-related macromolecular content (and secondarily to iron and tissue water fraction) (*30*); R_2_* to iron and to a lesser extent myelin (*31*); MTsat to macromolecular structures including myelin (*32*, *33*); and proton density (PrD) to tissue water fraction (*15*). For the SRT, we leveraged publicly available dorsolateral prefrontal cortex (DLPFC, Brodmann area 46/9) data generated using Visium (10x Genomics) (*34*). This dataset included 30 brain samples from 10 individuals without a neurological condition at the time of death (mean age = 49.51 years, 4/10 females).

Due to the availability of high quality SRT data, we focused this work on the human DLPFC. This is also a well defined area of the brain both for dissection and segmentation on MRI, and is involved in several neurological conditions, including neurodegenerative and neuropsychiatric disorders (*35–38*). We ensured that the DLPFC from both methods was partitioned into three regions, namely anterior, middle and posterior DLPFC, with the anatomical landmarks used during postmortem human tissue dissection to guide the DLPFC segmentation of MRI data. For the ultra-high resolution MRI data, the DLPFC was defined using structural MP2RAGE images brought into MPM space, and the cortex was segmented into four equivolume spatial domains (SpD_qMRI_) from superficial to deep grey matter (**Fig 1C-E**). We found that qMRI metrics varied approximately linearly over cortical SpD_qMRI_ i.e. across cortical depth, similar to previous reports (*39*, *40*) (**SupFig 1A**). For SRT data, we used a molecularly– and spatially-informed partitioning of SRT spots, with similar areas across the spatial domains (SpD_SRT_) but preserving the inherent cellular architecture of the DLPFC. Therefore, we defined twelve common regions of interest (ROIs) across four spatial domains and three DLPFC subregions spanning the anterior-posterior axis for both SRT and MRI data, with one-to-one matching between SpD_qMRI_ and SpD_SRT_ domains. Spatially, SpD_1_ maps to the intracortical region closest to the pial surface, whilst SpD_4_ maps to the intracortical region adjacent to the WM. In this way, we generated a spatial integration framework, which could be used to investigate the relationships between qMRI and SRT metrics.

### qMRI metrics significantly correlate with and predict glial cell counts

We first investigated the extent to which qMRI was associated with counts of specific cell types as predicted using SRT data (**Fig 2A**). We chose to focus our study on glial cell types, as their reactivity and spatial mobility makes them fast responding cell types in the brain, and are especially likely to be involved in early disease stages. Furthermore, glia can have high lipid and iron content, both of which are substantial determinants of qMRI values (*41*). Therefore, we wanted to explicitly evaluate whether there was enough sensitivity in qMRI to infer the presence of these cell types.

**Figure 2.**
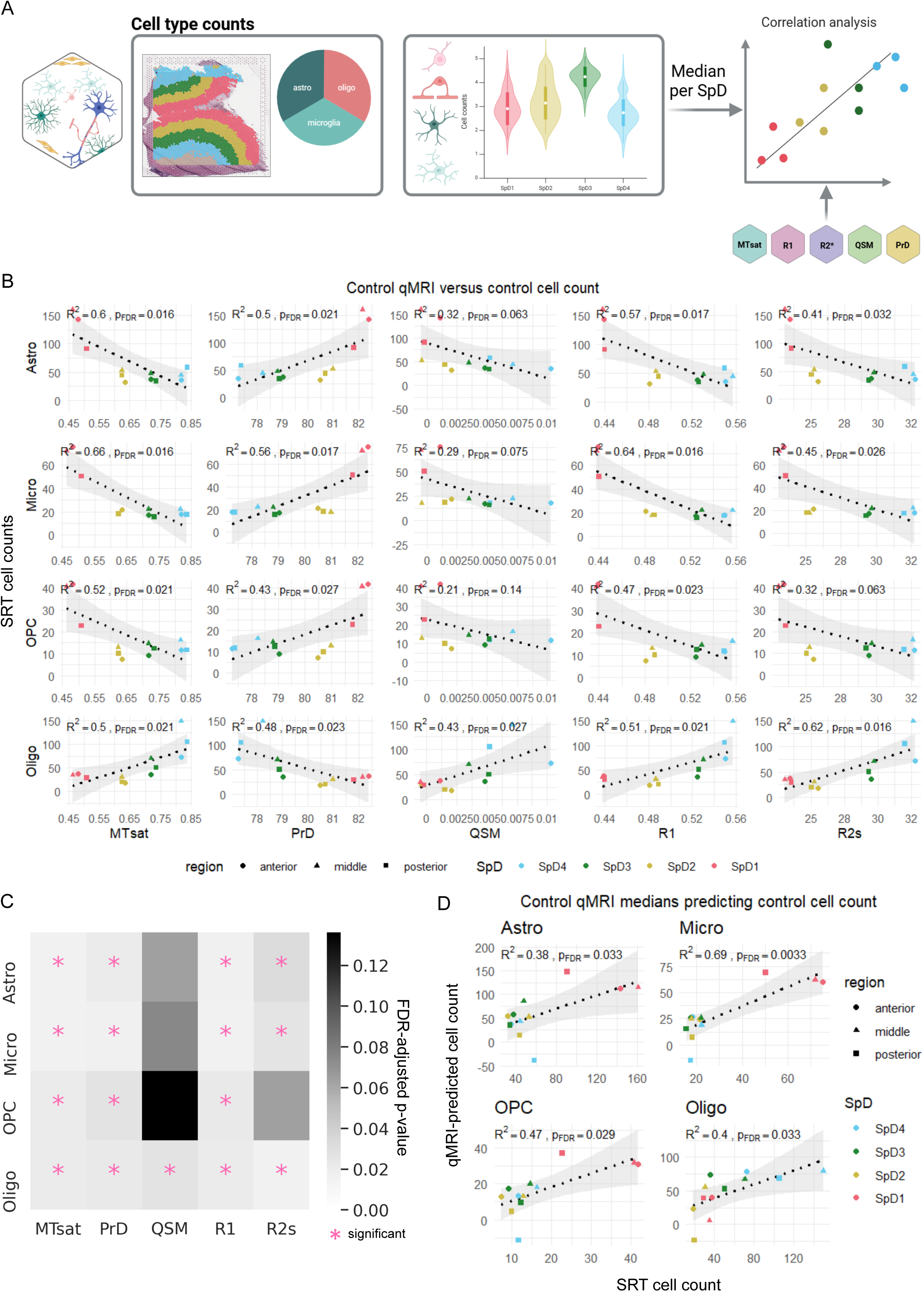
Characterising linear relationships between glial cell type proportions and qMRI metrics. (**A**) Schematic of cell type count data acquisition. SRT data were deconvolved using *cell2location*. Cell types with the top three *cell2location* scores were counted as present in a given spot. The median cell count per ROI was then correlated to the median qMRI metrics per ROI. (**B**) Correlative analysis of median SRT cell counts in relation to median qMRI metrics. (**C**) Heatmap summarizing FDR-adjusted p-values of correlations between cell counts and qMRI metrics. Significant pathways (where the FDR adjusted p-value<0.05) are indicated by a pink asterisk (*). (**D**) Correlation analysis of SRT predicted cell counts and qMRI predicted cell counts with LOOCV approach. All cell types have a significant correlation (FDR-adjusted p-value<0.05), although astrocytes, microglia and OPCs have a higher reliance on SpD_1_ ROIs for a linear fit.

The Visium technology used for the SRT data does not generate spatially resolved data at the single cell level, but rather each spatial unit (55 µm diameter spot) is estimated to contain 3-6 cells in human DLPFC (*34*). Therefore, to assess the relationship between cell type abundance and qMRI metrics, we first predicted which cell types were most likely to be in each given spatial unit. To do this, we used the deconvoluted cell types of this dataset (*34*), which were generated using *cell2location* (*42*). In brief, this approach used paired single nucleus RNA sequencing SRT data to predict the cell types most likely to be present in a given SRT spot (*34*, *43*). We considered the three cell types with the highest *cell2location* scores, since these were the ones for which we had highest confidence. This allowed us to sum the counts of each cell type across the entire ROI. This provided us with SRT-inferred cell counts for each glia type in each SpD_SRT_ across frontal, middle and posterior DLPFC in all samples (**SupFig 2A**). This, in turn, was summarised as the median SRT-inferred count across individuals, providing us with an estimated cell type count in each of the 12 ROIs. Meanwhile, for each SpD_qMRI_ we extracted median qMRI values within each ROI mask.

We opted for linear modelling to relate the SpD_qMRI_ and SpD_SRT_ data as a conservative, low complexity and robust strategy. We found that the cell type with the most significant relationships with qMRI metrics was oligodendrocytes, which had significant positive correlations with all qMRI metrics (**Fig 2B,C**, 0.43<R^2^<0.62, 1.60×10^−2^ < FDR-adjusted p-value < 2.66×10^−2^), apart from proton density, with which oligodendrocytes had a negative correlation (R^2^=0.48, FDR-adjusted p-value = 2.28×10^−2^). Given that oligodendrocytes have the highest myelin and iron content and these features dictate qMRI values for metrics with positive correlation, the observed correlations are biologically coherent. The strength and specificity of these associations provided new evidence linking cellular composition directly to *in vivo* MRI signatures. By contrast, microglia and astrocytes had significant negative correlations with all metrics except PrD with which they were positively correlated, and QSM with which they did not correlate. Due to their expected low myelin and lower iron content than oligodendrocytes, and the fact that their decrease from pial surface to white matter co-occurs with increasing oligodendrocyte density, this would be expected. A Cook’s distance analysis demonstrated that cell type count estimations were robust, as these were not sensitive to the removal of any one of the ROIs (**SupFig 2B**). Oligodendrocyte progenitor cells (OPCs) had the lowest number of significant correlations, having significant relationships with MTsat, PrD and R_1_ (**Fig 2C**). In summary, our correlation results are in accordance with what we would expect for each cell type, based on their myelin and iron content, as well as their known spatial distributions across the cortex.

We hypothesised that a multivariate qMRI model could be used to predict cell counts. Therefore, we aimed to verify whether cell type counts could be predicted using a linear combination of qMRI metrics. Predictive performance was assessed across 12 ROIs using leave-one-out cross validation (LOOCV) since this provided a robust framework for the number of samples available for this study (*44*). One ROI was iteratively withheld, while the multivariate model, based on a combination of qMRI metrics, was fitted across remaining ROIs and used to predict the cell type count derived by SRT. We found that the qMRI-predicted cell type counts significantly correlated with SRT counts for all cell types, with microglia having the strongest correlation (R^2^ = 0.69, FDR-adjusted p-value = 3.27×10^−3^, **Fig 2D**). Upon visual inspection, we found that the correlations for microglia, astrocytes and OPCs were largely reliant on the SpD_1_ scores, with low variation in the cell counts across SpD_2-4_. Therefore, to assess the robustness of the correlations in relation to SpD_1_, we calculated Cook’s distances for each ROI. We found that while specific ROIs impacted these cell types, SpD_1_ as a whole did not (**SupFig 2C**). In contrast, although oligodendrocytes (R^2^ = 0.40, FDR-adjusted p-value = 3.27×10^−2^) had a lower R^2^ and a higher FDR-adjusted p-value, they had a correlation which was less reliant on any individual SpD, making it more likely to be a robust relationship. As a result, we expect the prediction of microglia, astrocytes and OPCs cell type counts to be more sensitive to variations in SpD_1_, which is the spatial domain closest to the pial surface, but not as sensitive to deeper intra-cortical changes in SpD_2-4_. We also expect the cell type predictions of oligodendrocytes to be more robust.

### qMRI metrics correlate with and predict lipid, myelin and inflammation pathway scores

Having found strong associations between qMRI and cell counts, we next hypothesised that qMRI metrics would be associated with biological processes across matching SpD_qMRI_ and SpD_SRT_ pairs. In order to explore the molecular pathway-qMRI relationship, we chose to leverage gene ontology (GO) biological process (BP) pathways, which collate lists of genes involved in specific biological pathways (**Fig 3A**) (*45*). We chose to focus this investigation on four pathway categories or keywords: lipid, myelin, iron and inflammation. These were selected due to the expectation that qMRI metrics should capture features associated with these pathways. All pathways including these keywords were selected as well as all their offspring pathways (N_paths_ = 2,317). Pathways were then filtered based on our confidence to capture them on a molecular level, resulting in the exclusion of all pathways with five genes or less and all pathways where less than 50% of the genes were captured in our SRT dataset, resulting in a total of 1,270 pathways. This allowed us to score each spot with the expression of a pathway relative to background gene expression, defined as a list of genes of matching length, as previously described (*46*, *47*). For robustness, we bootstrapped this procedure and took as a final score the median value across bootstrapping iterations. Finally, we calculated the median GO pathway score of each pathway in each ROI (**SupFig 3**). As before, we extracted median qMRI values from the twelve ROIs. Similarly to the cell count estimation, we opted for linear modelling (**Fig 3B**).

**Figure 3.**
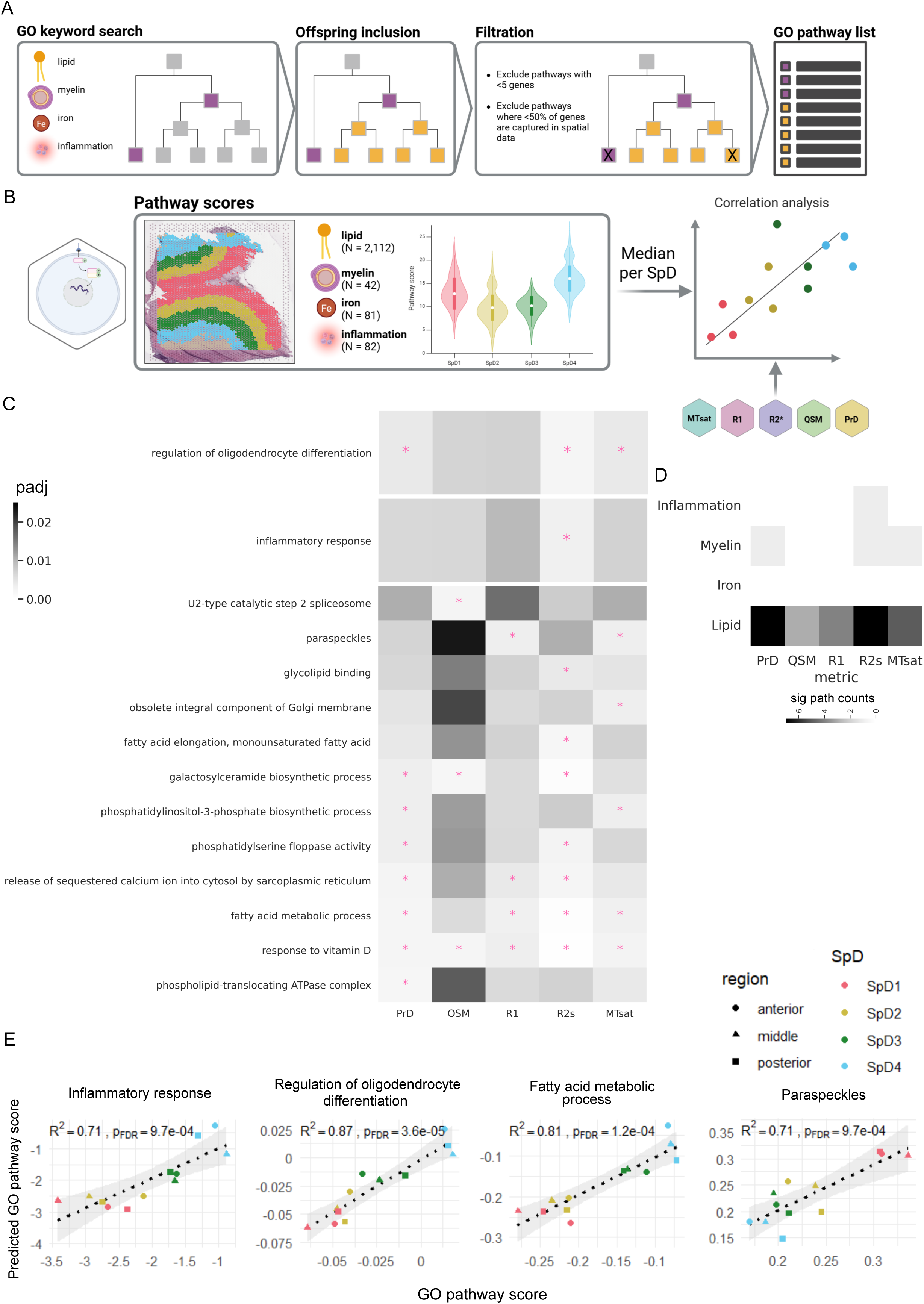
Characterising linear relationships between biological pathways and qMRI metrics. (**A**) Schematic of GO biological process pathway selection. All pathways containing selected keywords (lipid, myelin, iron, inflammation) and their offspring were included. All pathways which did not have at least 5 genes or which SRT data did not detect at least 50% of genes were excluded. (**B**) Pathway expression scores were calculated and correlative relationships with qMRI metrics were characterized. (**C**) Heatmap showing the FDR-adjusted p-value of all pathways with a significant correlation with at least one qMRI metric. Significant pathways (FDR adjusted p-value<0.05 and R^2^>0.85) are indicated by a pink asterisk (*). Terms are grouped in panels related to inflammation, myelin, and lipid metabolism. (**D**) Heatmap showing the sum of the number of pathways in each category significantly correlated with each qMRI metric. (**E**) Correlation of pathway scores predicted with LOOCV model based on qMRI data relative to real GO pathway scores from SRT data. We only show a small number of highlighted pathways.

A total of 14 pathways had scores which significantly correlated with qMRI metrics, where a pathway was considered to be significantly correlated if R^2^ > 0.85 and FDR-adjusted p-value < 5×10^−2^ (**Fig 3C, ST2**). Of these, one related to inflammation (inflammatory response, correlation with R_2_*), one related to myelin (regulation of oligodendrocyte differentiation, correlation with PrD, R_2_* and MTsat) and 12 related to lipid metabolism. PrD and R_2_* were the qMRI metrics with the highest numbers of significant correlations with specific pathway expression. These pathways related to various processes including splicing (U2-type catalytic step 2 spliceosome, paraspeckles), lipid metabolism (fatty acid elongation, galactosylceramide biosynthetic process, phosphatidylinositol-3-phosphate biosynthetic process, fatty acid metabolic process), transport and binding (phosphatidylserine floppase activity, release of sequestered calcium ion into cytosol by sarcoplasmic reticulum, phospholipid-translocating ATPase complex, glycolipid binding), and complex pathways such as response to vitamin D. Lipid pathways had the most significant correlations with qMRI metrics, and were mostly robust to excluding ROIs using Cook’s test (**Fig 3D**, **SupFig 4**).

Observing that many (5/14) GO BP pathways exhibited significant associations with at least three qMRI metrics, we modelled each GO pathway score as a linear combination of qMRI metrics using the same LOOCV approach across the 12 ROIs as described for cell counts. We found that for all of these pathways, the scores could be reliably predicted, with strong and highly significant correlations with the SRT scores (0.55<R^2^<0.92, 1.31×10^−5^< FDR-adjusted p-value < 5.77×10^−3^, **Fig 3E, SupFig 5A**). We then assessed the robustness of these correlations by testing the influence of individual ROIs on the correlations using Cook’s distances (**SupFig 5B)**. We found no consistent effects of ROI removal on pathway prediction. Overall, this suggests that the molecular scores for several biological processes, particularly lipid-associated processes, have significant and strong correlation relationships with qMRI data. Therefore, these processes are promising candidates to evaluate for non-invasive disease assessment.

### qMRI indicates disruption to lipids in living Parkinson’s disease patients

To assess the translatability of our approach, we applied our models to predict cell-type counts and pathway scores in a clinical cohort of living patients with PD (**Fig 4A**). PD follows a well-characterised anatomical progression, with subcortical regions such as the substantia nigra affected first and cortical regions later. This offers us a well established progression framework such that we can apply our method to a brain region expected to be largely unaffected anatomically, but which may show early signs of molecular changes.

**Figure 4.**
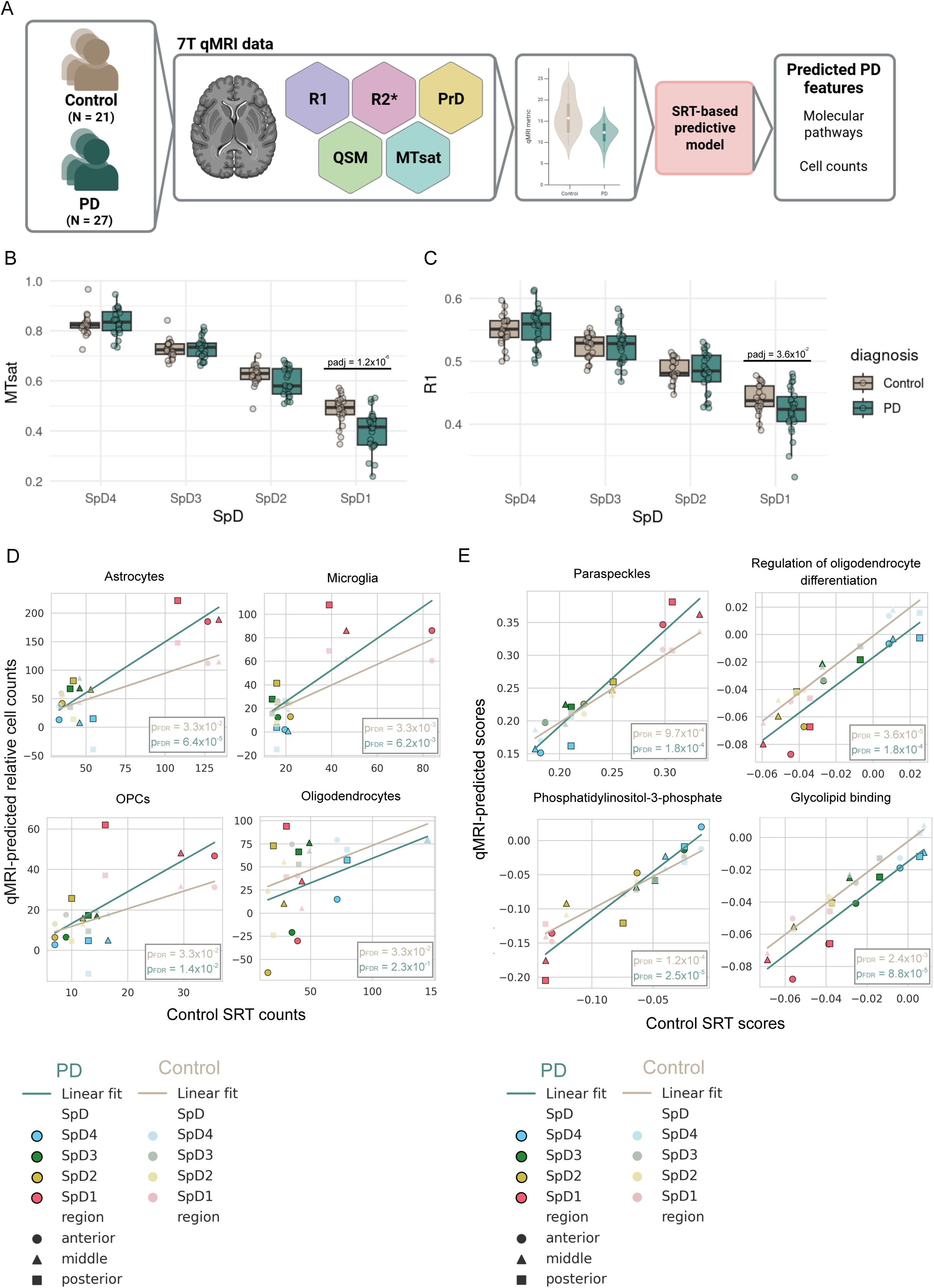
Prediction of molecular metrics in Parkinson’s disease patients. (**A**) Illustrative schematic of analysis strategy, comprising control and PD qMRI data. We applied the predictive model developed with control data to qMRI data collected from individuals with PD, predicting pathways and cell counts in disease. (**B**) MTsat levels across cortical spatial domains in cases and controls. We found a significant decrease in MTsat specific to SpD_1_ (FDR-adjusted p-value = 1.2×10^−6^). (**C**) R_1_ levels across cortical spatial domains in cases and controls. We found a significant decrease in R_1_ specific to SpD_1_ (FDR-adjusted p-value = 3.6×10^−2^). (**D**) Correlation analysis of predicted cell type counts in PD relative to control SRT deconvoluted counts. The linear model for astrocytes, microglia and OPCs have distinct beta in cases and controls (0.9< Z_β_ <2.0), whilst the model is maintained for oligodendrocytes (Z_β_ = 0.016). (**E**) Overlaid correlation of SRT score vs predicted pathway score for controls and PD.

We utilised 7T qMRI data from 27 people with Parkinson’s disease (PD, age = 67.3 ± 6.54 years, 6/27 females), acquired with the same protocol as for the cohort of healthy controls described above, and extracted qMRI metrics from the 12 ROIs using identical processing steps to the control cohort (**SupFig 6A,B**). First we assessed whether there were group differences in any qMRI metrics when examining the whole DLPFC, and found no significant differences for any (1.90×10^−1^ < FDR-adjusted p-value < 9.39×10^−1^, **SupFig 6C**). We then performed a pairwise comparison of control vs PD qMRI metrics across SpD_qMRI_s. We found a significant decrease in MTsat levels in PD, which was specific to SpD_1_ (FDR-adjusted p-value = 1.23×10^−6^, **Fig 4B**), along with a decrease in R_1_ in the same SpD_qMRI_ (FDR-adjusted p-value = 3.55×10^−2^, **Fig 4C**). This demonstrated to us the value of our approach using spatial partitioning of qMRI data within brain regions.

To investigate the biological basis of the SpD_1_-specific changes to MTsat and R_1_, we applied the SRT-based model to generate cellular and molecular predictions in PD (**ST3**). As in controls, we obtained qMRI-based cell count predictions for PD, enabling direct comparison with qMRI-predicted control counts, using the SRT-predicted control counts as the reference. We evaluated these relationships in three stages. First, we determined whether the qMRI-predicted PD values remained significantly correlated with the SRT-predicted control counts, as observed previously in controls. Loss of correlation would indicate deviation from the control molecular architecture of the grey matter cortex in PD. Second, we compared the slopes as indicated by the beta coefficients of the PD and control models to determine whether the rate of variation across the cortical grey matter differed in PD as compared to control groups. Third, in cases where the slopes (beta coefficients) were similar, we assessed differences in the intercepts, with the aim of determining whether there were uniform upward or downward shifts in predicted counts across all SpDs. For these analyses, we compared two linear models: qMRI-predicted counts in controls regressed on SRT-predicted control counts, and qMRI-predicted counts in PD regressed on the same SRT-predicted values (**ST4**). We computed Z-tests for slope (β) and intercept differences and used ordinary least squares regression to determine whether these differences were associated with PD status.

We found that for astrocytes, microglia, and OPCs, correlations between qMRI-predicted cell counts and SRT-predicted control counts remained significant in PD (**Fig 4D**). The slope differed between groups, with higher beta slope coefficients in PD than in controls (Astrocytes: Z_β_ = 2.09; Microglia: Z_β_ = 0.99; OPCs: Z_β_ = 1.06), largely driven by increased qMRI-predicted counts in PD SpD_1_. Associations with PD were weak (1.96×10^−2^ < FDR-adjusted p-value < 4.40×10^−2^), and only astrocytes reached nominal significance in the association with PD (nominally significant, p-value = 4.90×10^−2^).

For oligodendrocytes, the correlation was no longer significant (R² = 0.137; FDR-adjusted p-value = 2.35×10^−1^), although the slope remained similar between PD and controls (Z_β_ = 1.60×10^−2^; **Fig 4D**). Predicted oligodendrocyte counts were generally lower in PD, reflected in a minimal beta change and a negative intercept shift (Z_INT_ = –0.50). Together, these findings suggest reduced predicted oligodendrocyte counts in PD that vary across cortical SpDs. Large negative predicted counts in some ROIs indicate that PD tissue has lower oligodendrocyte abundance than the control data used to train the model.

Using the LOOCV approach, we next compared predicted pathway scores between PD and control groups, anchoring both to SRT control pathway scores. Three pathways showed significantly lower predicted scores in PD, namely oligodendrocyte differentiation (FDR-adjusted p-value = 4.01×10^−2^), glycolipid binding (FDR-adjusted p-value = 3.75×10^−2^), and the U2-type catalytic step 2 spliceosome (FDR-adjusted p-value = 3.54×10^−2^; **SupFig 7A**). These findings paralleled our cell-count analyses and point toward coordinated alterations in oligodendrocyte-and lipid-related processes in PD.

All PD pathway predictions remained significantly correlated with SRT pathway scores (**Fig 4E**, **SupFig 7B**), though several pathways showed altered beta slope coefficients (–1.52 < Z_β_ < 2.85, **SupFig 8**). Two pathways, phosphatidylinositol-3-phosphate (nominally significant, p-value = 2.76 × 10^−2^) and paraspeckles (nominally significant, p-value = 9.67×10^−3^), showed nominally significant beta shifts, suggesting changes in their spatial gradients across the PD cortex. For glycolipid binding and oligodendrocyte differentiation, the pathways with the clearest overall reductions, the slopes were largely unchanged (glycolipid binding: Z_β_ = 2.98×10^−2^ and oligodendrocyte differentiation: Z_β_ = –1.46×10^−1^), but both showed negative intercept shifts (glycolipid binding: nominally significant, p-value = 4.35×10^−2^, Z_INT_ = –2.15; oligodendrocyte differentiation: nominally significant, p-value = 2.32×10^−2^, Z_INT_ = –2.45). This pattern indicates a global downward shift in pathway activity rather than a redistribution across cortical depth.

Together with the reduced oligodendrocyte predictions, these findings suggest broad impairment of lipid-associated and oligodendrocyte-related biology in PD grey matter, consistent with emerging evidence of disrupted myelin and lipid homeostasis in the disease.

### Transcriptomics corroborate qMRI-predicted glial dysregulation in Parkinson’s disease

The qMRI-based predictions highlighted oligodendrocytes and astrocytes as the two cell types most likely to show altered proportions in PD. For oligodendrocytes, both the cell-count predictions and pathway analyses suggested disruptions consistent with reduced oligodendrocyte abundance and impaired maturation. We therefore assessed whether these predictions were supported by independent transcriptomic data. To do so, we used previously generated snRNA-seq data from mid-stage PD (Braak 3/4) (*48*) to match the disease stage of the living PD cohort in our qMRI dataset. This dataset includes samples from the frontal, cingulate and parietal cortex regions from controls (N_controls_ = 20) and PD cases (NPD = 18), with comparable age and sex distributions (*48*). All major CNS cell types were recovered following quality control and preprocessing, including neurons, oligodendrocyte-lineage cells, astrocytes, microglia, immune cells, and endothelial/mural cells.

Consistent with the qMRI predictions, oligodendrocyte maturation genes were significantly enriched among downregulated genes in frontal cortex astrocytes (FDR-adjusted p-value = 3.09×10^−3^, **Fig 5A**). The enrichment within astrocytes aligns with their known role in supporting CNS myelination (*49*, *50*) and suggests that this interaction may be perturbed in PD. Bulk RNA-seq data provided additional support (**SupFig 9A**). Given these transcriptional signatures of reduced maturation capacity, we next examined whether oligodendrocyte-lineage proportions were diminished in PD. Indeed, cell-type proportion analysis revealed a significant reduction in oligodendroglial cells (logFC = –0.197, FDR-adjusted p-value = 4.05×10^−2^, **Fig 5B,C**, **ST5**), directly supporting the qMRI-based predictions.

**Figure 5.**
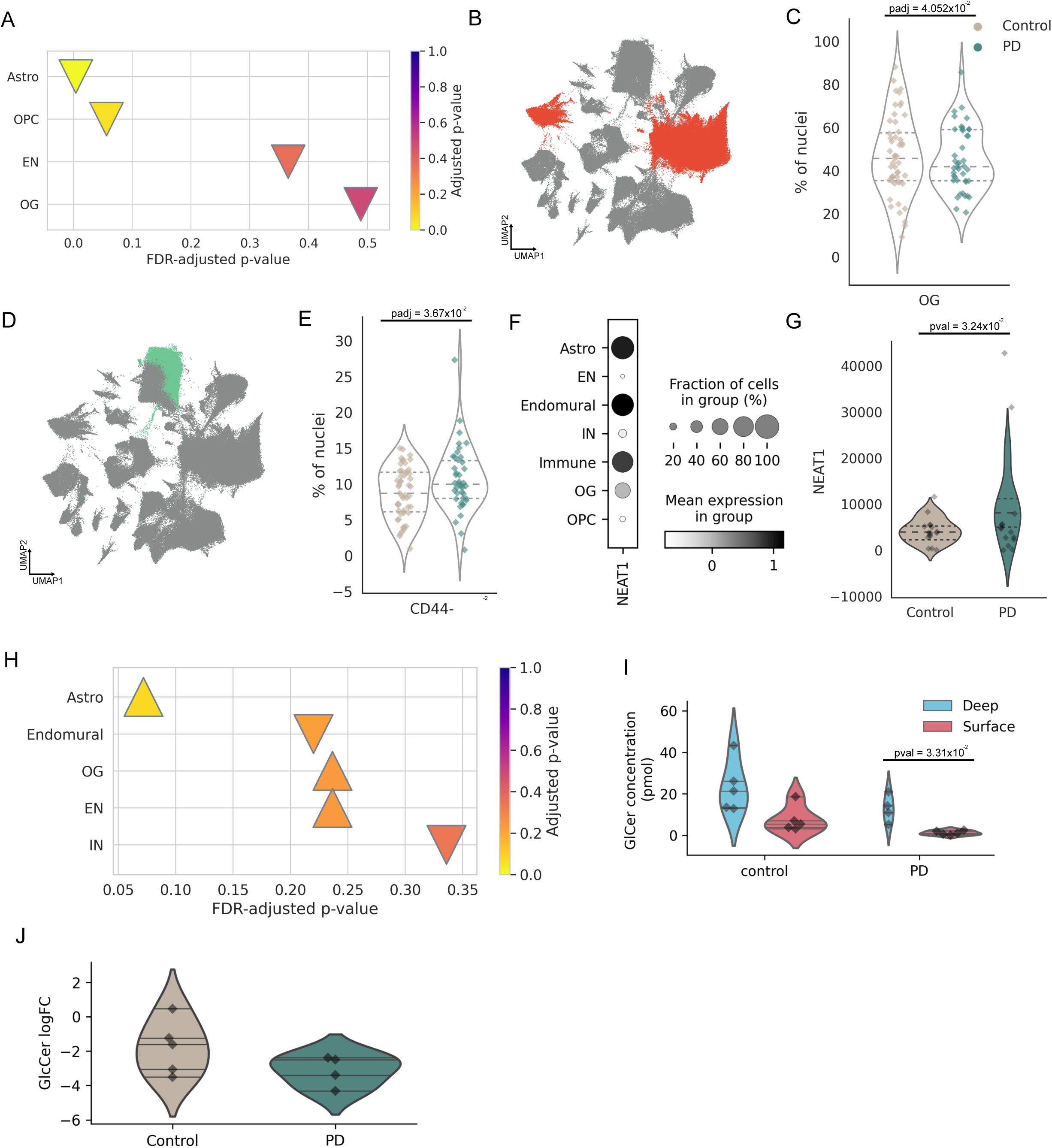
Predicted changes to oligodendrocyte population in PD can be validated through other molecular methods. (**A**) Fisher enrichment test of oligodendrocyte maturation genes in mid-stage (Braak 3/4) PD snRNA-seq dataset. This gene set is enriched in down-regulated genes in astrocytes and have nominally significant enrichment in down-regulated genes in OPCs. (**B**) UMAP of single nucleus RNA sequencing data from 740,000 nuclei from mid-stage (Braak 3/4) PD (*48*). Cells of the oligodendrocyte progenitor lineage are highlighted in orange. (**C**) Cell type proportion analysis of cells of the oligodendrocyte progenitor lineage in PD compared to controls, with a significant decrease in disease (logFC = –0.197, FDR-adjusted p-value = 4.0×10^−2^). (**D**) UMAP of single nucleus RNA sequencing data highlighting CD44^−^ protoplasmic astrocytes. (**E**) Cell type proportion analysis comparing the proportion of CD44^−^ astrocytes in PD and controls. (**F**) Dotplot showing mean expression levels of *NEAT1* across major cell types, where the dot size represents the % of the cell type population with positive expression of that gene. Astrocytes, the endomural compartment and immune cells have the highest *NEAT1* expression. (**G**) RNA-seq differential gene expression of *NEAT1* in the frontal cortex protoplasmic astrocyte population, resident to grey matter (nominally significant, logFC = 0.489, FDR-adjusted p-value = 3.24×10^−2^). (**H**) Fisher’s enrichment test of phosphatidylinositol-3-phosphate biosynthetic process pathway genes among differentially expressed genes in the frontal cortex, when comparing PD to controls. (**I**) HPLC quantification of GlcCer in superficial and deep cortex from controls and PD. There is a significant difference in GlcCer in the surface compared to deep cortex in PD (p-value = 3.31×10^−2^), which is not present in controls. (**J**) Violin plots showing log_2_FC in the surface cortex relative to deep cortex (Mean_Control_ log_2_FC = –1.788907; Mean_PD_ log_2_FC = –3.147041).

We then assessed astrocytes, focusing on the CD44^−^ population (**Fig 5D**), which corresponds to protoplasmic astrocytes enriched in grey matter, the compartment captured by our qMRI-based modelling. We found a significant increase in CD44^−^ astrocytes in PD (logFC = 0.093, FDR-adjusted p-value = 3.67×10^−2^, **Fig 5E**). This direction of effect mirrors the qMRI-predicted elevation of astrocyte signal in SpD_1_, the superficial cortical layer closest to the pia, where CD44^−^ astrocytes are most abundant. Notably, this signal was not detectable when analysing total astrocyte proportions, underscoring the added value of a spatially-informed approach (**SupFig 9B**).

Finally, we examined whether transcriptomic data supported qMRI-predicted disruptions in the paraspeckles and phosphatidylinositol-3-phosphate pathways. For paraspeckles, we assessed *NEAT1*, the essential long non-coding RNA scaffold of paraspeckle structures (*51*). The highest qMRI-predicted pathway activity occurred in SpD_1_, and correspondingly *NEAT1* expression was highest in astrocytes, immune cells, and endothelial cells (**Fig 5F**), which are cell types enriched in this superficial layer (**SupFig 3A**). Of these major cell types, a specific subpopulation of CD44^−^ cortical astrocytes (logFC = 0.489, FDR-adjusted p-value = 3.24×10^−2^; **Fig 5G**) and cycling macrophages (logFC = 1.35, FDR-adjusted p-value = 4.20×10^−2^) had nominally significant *NEAT1* upregulation, consistent with a spatially specific pathway shift. This was also supported by bulk RNA-seq (**SupFig 9C**). We also found transcriptomic evidence supporting altered phosphatidylinositol-3-phosphate biology, with nominal enrichment of upregulated genes in frontal cortex astrocytes (nominally significant, p-value = 9.38×10^−3^; **Fig 5H**). Together, these transcriptomic analyses independently confirm the key qMRI-derived predictions: reduced oligodendrocyte maturation and abundance, altered astrocyte populations with spatial specificity, and disruptions in paraspeckle– and PI3P-related pathways. These convergent results demonstrate that qMRI can capture biologically meaningful, cell-type-specific molecular changes in living PD patients.

### HPLC confirms superficial cortex-specific glycolipid reductions in Parkinson’s disease

We next assessed whether glycolipid alterations could support the qMRI prediction of reduced glycolipid content in PD. Glycolipids (GLs), a major class of CNS lipids composed of a hydrophobic ceramide backbone linked to a glycan headgroup, exist in numerous forms that vary in carbohydrate structure, chain length, and ceramide saturation or hydroxylation. GLs contribute to diverse neural functions, including cell differentiation, cell–cell interactions, dendritic development, synaptic formation and transmission, and energy metabolism. Since the GO glycolipid binding pathway reflects interactions between glycolipids and partner molecules, reductions in GL abundance would be expected to diminish pathway activity.

We first examined this transcriptomically by testing whether GL-associated genes were enriched among differentially expressed genes across cell types. Using an in-house glycosphingolipid (GSL) gene set, we found no significant enrichment (**ST6**). We assessed gene expression of *UGCG*, the enzyme synthesising glucosylceramide (GlcCer) which is a precursor of many GSLs, but it was not significantly differentially expressed in any cell type (non-significant, 5.81×10^−2^< p-value < 9.53×10^−1^). These observations suggested that glycolipid alterations in PD may be spatially localised or regulated post-transcriptionally, and therefore not captured by global transcriptional analyses.

Guided by the qMRI prediction that glycolipid binding pathway score was especially reduced in SpD_1_ (**Fig 4E**), we hypothesised that GL differences might be cortical-depth-specific. To test this, we quantified GSL and GlcCer concentrations in superficial and deep cortex from postmortem PD and control brains (N_controls_ = 5, N_PD_ = 5) using high-performance liquid chromatography (HPLC). Since intracortical variation was of particular interest and paired comparisons improve statistical power, we performed paired analyses within individuals. GlcCer concentrations were significantly lower in the superficial cortex relative to the deep cortex in PD (logFC = –3.147, p-value = 3.31×10^−2^), whereas this difference was not significant in controls (logFC = –1.788, p-value = 1.02×10^−1^) (**Fig 5I,J, ST7**). For glycosphingolipids, only a nominally significant difference in Gb3 was observed (pval = 1.99×10^−2^, FDR-adjusted p-value = 3.78×10^−1^, **SupFig 10**). As the glycolipid binding pathway depends on interactions involving GLs, reduced GL concentrations would be expected to lower binding activity, providing molecular support for the qMRI-predicted disruption of glycolipid pathways in PD.

In summary, our SRT-informed qMRI model inferred changes in cell types and molecular pathways in the living human brain, including alterations in oligodendrocytes, paraspeckle biology, and glycolipid pathways. These predictions were independently validated across bulk RNA-seq, snRNA-seq, and HPLC analyses, with consistent directionality and spatial specificity. Together, these results highlight the utility of our approach for non-invasive prediction of molecular and cellular features in living patients.

## Discussion

The emergence of disease modifying therapies for neurodegenerative disorders has intensified the need for non-invasive tools capable of capturing dynamic molecular pathology in living patients. Although qMRI is widely accessible and clinically established, its molecular interpretability has been limited. Spatially resolved transcriptomics (SRT), by contrast, provides spatially anchored molecular information that can be used to map MRI contrast to specific cellular and molecular processes. Here, we demonstrate the first SRT-informed modelling of qMRI at laminar resolution in the human cortex, enabling inference of molecular and cellular features from qMRI data acquired in vivo.

Spatial subdivision is a foundational principle in modern molecular neuroscience, where methods such as SRT and single nucleus RNA-seq reveal cell type– and pathway-level structure that bulk approaches obscure. As ultra-high resolution qMRI increasingly captures microstructural variation across cortical depth, an analogous opportunity arises in neuroimaging. Integrating qMRI with SRT across cortical laminae allows biological signals embedded within grey-matter microstructure to be resolved, moving beyond region-level associations (*52–55*) and providing a framework for interpreting qMRI metrics in terms of specific cellular and molecular entities.

Applying this framework in neurologically healthy brains revealed multiple significant correlations between glial cell counts and qMRI metrics. This corroborates previous findings showing strong correlations between qMRI metrics and von-Economo cell counts across cortical depths, as well as between qMRI and layer-specific genes in the Allen Human Brain Atlas (*40*). Oligodendrocytes were the cell type most strongly predicted by qMRI metrics, reflecting qMRI’s sensitivity to myelination (*56*). qMRI signals also correlated with pathways related to lipid metabolism, myelin and inflammation, with lipid-associated pathways showing the strongest and most consistent associations. These findings suggest that qMRI is particularly sensitive to lipid-rich microstructural environments and can capture biologically meaningful variation within the grey matter.

When we applied our model to Parkinson’s disease (PD), it generated a coherent set of predictions pointing to disruptions in oligodendrocyte biology, paraspeckle function, the phosphatidylinositol-3-phosphate (PI3P) biosynthetic pathway and glycolipid binding. These predictions provided a biologically interpretable scaffold for evaluating whether qMRI could detect molecular features known or hypothesised to be altered in PD. For example, a major predicted alteration was a reduction in oligodendrocyte abundance and impaired differentiation. Both single-nucleus and bulk RNA-seq supported this, revealing reduced maturation of the oligodendrocyte lineage, downregulation of maturation markers and lower proportions of oligodendroglial cells in PD cortex. These observations align with genetic evidence implicating oligodendrocytes in PD risk (*57*),(*58*).

The model also predicted disruption of the PI3P pathway, driven primarily by astrocytic changes. PI3P dysregulation promotes α-synuclein aggregation, and astrocytic responses to α-synuclein can alter myelination and lipid homeostasis (*59*), providing a mechanistic link between glial activation and impaired myelin-related processes (*49*, *60*, *61*). Predicted reductions in glycolipid (GL) activity were validated biochemically: high-performance liquid chromatography revealed a significant decrease in glucosylceramide specifically in the superficial cortex of PD donors, mirroring the spatial pattern predicted from qMRI. This is consistent with the established role of GBA1 and glycolipid metabolism in PD progression and cognitive decline (*62*, *63*), and suggests that our approach may be sensitive to early lipid disruptions relevant to clinical outcomes.

Beyond glial and lipid pathways, the model predicted disruption of paraspeckle biology, with the strongest effects in superficial cortical layers. We confirmed upregulation of *NEAT1*, which is the scaffold lncRNA essential for paraspeckle formation (*64*), particularly in protoplasmic astrocytes-enriched layers. Paraspeckle dysfunction has been linked to cognitive impairment in neurodegenerative models (*65*), raising the possibility that this molecular signature has prognostic value in PD. The ability of qMRI-derived features to capture such nuclear regulatory processes highlights the broader potential of SRT-guided models to extend qMRI’s interpretability beyond traditional structural or myelin-related domains.

This proof-of-concept study has several limitations. Both qMRI and SRT datasets were modest in size and the prediction model was trained exclusively on control SRT data. Although 0.6 mm resolution does not fully resolve individual cortical layers, our laminar partitioning nonetheless revealed spatially informative molecular associations. With the emerging availability of 11.4T systems, SpD partitioning could match the more biologically– and neuronally-relevant cortical layer structure. Differences between living and postmortem tissue (*66–68*), particularly regarding inflammatory states, may weaken some signals; however, oligodendrocyte, lipid and myelin-related features appear robust (*69*). Larger multimodal datasets, including disease SRT, will strengthen model generalisability, particularly for machine-learning approaches which may uncover nonlinear relationships between qMRI and SRT features. Finally, SRT-informed modelling may guide the development of new qMRI sequences optimised to detect specific molecular signatures.

Taken together, our findings show that SRT-informed qMRI provides a scalable, tolerable and non-invasive platform for inferring molecular and cellular features in the living human brain. Validated across transcriptomic and biochemical modalities, this approach reveals glial, lipid and nuclear regulatory processes relevant to PD, and has the potential to transform mechanistic research, patient stratification and clinical trial design across neurodegenerative diseases.

## Code availability

All code associated with this project is publicly available at https://github.com/mgrantpeters/MRIxST. All bulk and single nucleus RNA sequencing data will be made available through the CRN cloud (https://cloud.parkinsonsroadmap.org/collections) and is available upon request in the meantime.

## Acknowledgments

MGP is supported by the MS society (project number RRAG/480). AZ is supported by an Alzheimer’s Research UK Clinical Research Fellowship (2021B-001) and received grant funding from the Academy of Medical Sciences, Rosetrees and Parkinson’s UK. RW is supported by a Wellcome Career Development Award (225263/Z/22/Z) and has received grant funding from the UCLH Biomedical Research Centre, Parkinson’s UK (G-2404), the Lewy Body Society, and Rosetrees Trust. KS is supported by a European Research Council Consolidator Grant (DiSCo MRI SFN 770939). This work is supported by the UK Dementia Research Institute (award number UK DRI-2206) through UK DRI Ltd, principally funded by the Medical Research Council (MR) and the Lieber Institute for Brain Development. We gratefully thank the families who donated tissue to make this research possible. We thank the families of Connie and Stephen Lieber and Milton and Tamar Maltz for their generous support of this work.

## Materials and Methods

### MRI participants

We recruited 21 healthy controls aged 50 to 80 years from sources including university volunteer databases. Exclusion criteria were presence of neurological or psychiatric disorders, dementia, or contraindications to MRI. 32 age-matched people with PD within 10 years of diagnosis were also recruited to our London centre. Inclusion criteria were clinically diagnosed, early to mid-stage PD (Queen Square Brain Bank Criteria). Exclusion criteria were confounding neurological or psychiatric disorders, dementia, or contraindications to MRI. Participants continued their usual therapy (including levodopa) for all assessments. No patients were taking cholinesterase inhibitors. All participants gave written informed consent and the study was approved by the local Research Ethics Committee. In addition, After MRI quality control (conducted blinded to patient group), five PD participants were excluded due to excessive motion artefacts. Thus, 27 participants with PD (age 66.2 ± 7.24 years, mean ± SD; 6 female) and 21 healthy controls (age 67.7 ± 7.96 years, mean ± SD; 10 female) were included in the current study (**ST1**).

### MRI acquisition

All participants underwent multiparametric mapping (MPM) as well as 3D magnetization-prepared two-readout rapid acquisition gradient-echo (MP2RAGE) anatomical imaging on a 7T MRI system (Magnetom Terra, Siemens Healthineers, Erlangen, Germany) equipped with an eight-channel transmit/32-channel receive radio-frequency head coil (Nova Medical, Wilmington, MA) operating in a quadrature-like “TrueForm” mode. The MPM protocol consisted of three multi-echo fast low angle shot (FLASH) scans covering the whole brain with T1, proton density and magnetization-transfer weighting (T1w, PrDw and MTw) respectively that had 0.6 mm isotropic resolution, a readout bandwidth of 469 Hz/pixel and an acquisition time of 9 min 10 s each enabled by an overall parallel imaging acceleration factor of 4. The T1w scan was acquired with a flip angle (FA) of 24°, six evenly spaced echoes of alternating polarity at echo times between 2.3 and 14.2 ms and a TR of 19.5 ms; the PrDw scan with an FA of 6°, six evenly spaced echoes of alternating polarity at echo times between 2.2 and 14.1 ms and a TR of 19.5 ms; and the MTw scan with an FA of 6°, four evenly spaced echoes of alternating polarity at echo times between 2.2 and 9.34 ms, a TR of 19.5 ms and an MT contrast inducing off-resonant Gaussian pre-pulse with a flip angle of 180°. Before each sequence, a low resolution 4mm isotropic image covering the same field of view was acquired to measure the relative radio-frequency receive coil sensitivity between contrast weightings (*70*). An estimate of the radio-frequency B_1_^+^ transmit field was calculated at the time of scanning using a 3D Bloch-Siegert based approach (*71*). Magnitude and phase images for each of the MPM contrasts, as well as the B_1_^+^ maps, were reconstructed using MORSE, a real-time image reconstruction pipeline (*72*). Whole-brain MP2RAGE scans were acquired with the following parameters: Field of view = 14.4 x 20.4 x 20.4 cm, 0.65 mm isotropic resolution, TR = 5000 ms, TE = 2.54 ms, TI1 = 900 ms, TI2 = 2750 ms, FA1 = 5°, FA2 = 3°, readout bandwidth = 240 Hz / pixel and an acquisition time of 8 min 42 s.

### qMRI map creation

Quantitative MPM maps of the longitudinal relaxation rate (R_1_), transverse relaxation rate (R_2_*), proton density (PrD), and magnetization transfer saturation (MTsat) were calculated using the hMRI toolbox v0.5.0 (*73*) for Statistical Parametric Mapping (SPM) (https://www.fil.ion.ucl.ac.uk/spm/software/spm12/) in MATLAB (R2018a, The MathWorks Inc., Natick, Massachusetts, USA). All acquisitions were coregistered to their corresponding PrDw scan using SPM12. R_1_ maps were estimated from the PrDw and T1w data using approximations of the signal equations for small flip angle (*74*), (*75*). Correction for imperfect spoiling was completed using local scanner-sequence acquisition-specific parameters (*76*). R_2_* maps were derived from the mono-exponential TE dependence of the PrDw, T1w and MTw signals using the ESTATICS model and weighted least squares fitting (*77*). MTsat maps were calculated from the MTw sequence, accounting for spatially varying R_1_ relaxation and transmit field efficiency (*74*). Position-dependent receive field variation was corrected using the “per contrast” option and the low resolution calibration pre-scans. B_1_^+^ bias correction was completed using the pre-calculated map from the Bloch-Siegert acquisition.

Quantitative susceptibility mapping (QSM) maps were generated using the MPM data by applying the following processing pipeline to PrDw and T1w data: 3D complex data were separated into odd and even echo trains and non-linear fitting of the complex data over echoes (*78*) was applied to each echo train before unwrapping the fitted field maps with a rapid path-based minimum spanning tree algorithm (ROMEO) (*79*). Unwrapped field maps from odd and even echo trains were scaled to Hz and averaged to create a single field-map. To increase QSM map accuracy, field maps were rotated to align the inferior-superior axis with the direction of the scanner’s main magnetic field (*80*). Removal of background fields was completed using projection onto dipole fields (PDF) (*81*), and brain masks for this step were generated by thresholding the phase quality maps generated during echo fitting. Dipole inversion was completed using STAR-QSM (*82*). Susceptibility maps created from T1w data were coregistered to their corresponding PrDw space using SPM12 before both maps were averaged to give a final QSM map (*83*), (*84*).

### qMRI region of interest (SpD_qMRI_) extraction

Prior to structural processing, T1w MP2RAGE (UNI) images were co-registered to their corresponding PrDw MPM space using SPM12. These coregistered T1w images were then processed using the ‘recon-all’ pipeline with the ‘highres’ setting in Freesurfer version 7.1.0 (*85*). Errors in cortical surface reconstruction were manually corrected after visual inspection. The grey and white matter surfaces generated by Freesurfer were then used to define 12 equi-volume cortical depth volumes using LayNii (*86*), which were grouped together to produce the final four depth volumes or spatial domains SpD_1-4_ (*87*). A version of the HCP-MMP1.0 atlas (*88*) projected onto fsaverage space was then brought into each subject’s native space using surface-based registration in Freesurfer. Brodmann area 46 as defined in the HCP-MMP1.0 atlas was used as the DLPFC label, and this was transformed into a volumetric mask extending from the pial surface to the grey / white matter boundary in PrDw MPM space using Freesurfer. Using custom MATLAB scripts, the DLPFC was further divided into anterior, middle and posterior equi-volume sub-regions, before being intersected with the four depth-volume masks to create 4 x 3 individual ROIs for each subject. Median QSM, R_1_, R_2_*, PrD, and MTsat values were extracted from each of these ROIs using MATLAB.

### SRT dataset description

Publicly available SRT data from postmortem DLPFC was used for this analysis (*34*). Briefly, the sample set consists of 30 tissue blocks from 10 donors. For each donor, anterior, middle and posterior DLPFC samples were dissected and profiled. Brodmann area 46 was dissected in a plane perpendicular to the pial surface to capture cortical layers 1-6 and WM. The 10x Genomics Visium protocol was performed according to manufacturer’s instructions on 10 um sections. Sequencing was performed on an Illumina NovaSeq 6000 with a minimum depth of 50,000 reads per Visium spot as previously described (*34*).

### SRT layer (SpD_SRT_) partitioning

Data were used following processing previously described (*34*). Briefly, FASTQ and image data were pre-processed with the 10x Genomics SpaceRanger pipeline. Outputs from this step were processed in R, including filtration of undetected genes and spots with zero counts and QC metric assessment. This was followed by dimensionality reduction (n=50 PCs) and batch-correction using Harmony (*89*). Clustering was performed in a data-driven and spatially-aware manner using BayesSpace (*90*).

We opted to partition the spatial domains of the SRT data with a compromise of molecular feature and layer size compatible with the qMRI partitioning. To do this, we used BayesSpace clustering with n=7 clusters which partitioned the cortex into 5 spatial domains, of which cluster 1 consisted of a small endothelial-rich transcriptomic signature. Aware that this cluster would be included as part of SpD1 in the MRI data, we combined this cluster manually with cluster 2, resulting in 4 cortical spatial domains.

### SRT SpD mapping to molecular cortical layers

Spatial registration pipeline from *spatialLIBD* v1.21.5 was used to relate the 5 SRT SpDs to the classical layer of the cortex (*91*). Manual annotation of six grey matter layers and white matter in SRT DLPFC data from Maynard et al. was used as a reference (*92*). Analysis was completed with tools from *spatialLIBD* v1.23.0: enrichment stats for 5 SRT SpDs were calculated with `registration_wrapper(covars = c(“position”, “age”, “sex”))`, modeling data from Maynard et al (*92*) were accessed with `fetch_data(type = “modeling_results”)`, SpD correlations calculated with `layer_stat_cor()`, and visualised with `layer_stat_cor_plot()`.

### Gene ontology pathway selection

Existing literature predicts that qMRI metrics relate to specific biological processes. Keywords associated with these processes – namely, myelin, iron, mineral and lipids – were searched among gene ontology biological process pathways (https://geneontology.org/) (*45*). A list of pathways associated with these keywords was downloaded on 2024-11-11. Due to the hierarchical nature of GO pathways, we opted to also include all offspring pathways of our pathways of interest, since this could provide further insight into specific processes, for which we used the *GO.db* R package v3.16.0. This provided us with a list of all molecular pathways and their offspring we expected to be associated with entities captured in qMRI data. We then used the R package *AnnotationDbi* v1.60.2 to fetch all gene symbols associated with each of the pathways of interest.

### Spatial scoring of GO molecular pathways

In order to determine the level of expression of our pathways of interest, we adapted the *scanpy* (*47*) `score_genes` function to take in a custom gene background list. In brief, this function sums the expression of a list of genes of interest. It then subtracts from this score the summed expression of a list of randomly selected background genes of matching length. We made two modifications to this approach, firstly by allowing the use of a custom background gene list. This allowed us to use a gene ‘universe’ of genes captured in the data minus the genes of interest to avoid double-sampling. Secondly, following the subtraction of the background score, we divided our score by the sum of UMI in that SRT spot. This was done to account for layer-specific differences in UMI capture due to the differences in tissue permeabilisation in the white and grey matter, making scores more comparable across spatial domains regardless of myelination state.

### SRT layer-specific cell count estimation

The spatialDLPFC dataset was deconvoluted using *cell2location* (*42*) with paired single nuclear RNA sequencing data (*34*). This assigns to each spot a likelihood score of a given cell type being present within that spot. In order to ensure only the cell types we had the highest confidence of being present were counted, we considered the top 3 cell types with highest score to be present in a given spot.

### Correlation analyses

qMRI metrics, cell count and GO pathway scores were adjusted for age and sex across ROIs using the covariance method (*93*). Median, adjusted qMRI metrics, cell counts and GO pathway scores were calculated across subjects for each of the 4 x 3 ROIs. The Spearman correlation coefficients were then calculated for median qMRI metrics versus median cell counts, and median qMRI metrics versus median GO pathway scores. Significant correlations were presented at FDR-corrected p<0.05 (*94*).

### Prediction of control SRT from qMRI data

Leave-one out cross validation (LOOCV) was completed by fitting a linear regression model to all but one ROI at a time like so: cell count or GO pathway score ∼ QSM + R_1_ + R_2_^*^ `+ MTsat + PrD.` The cell count or GO pathway score for the left out region was predicted using the coefficients from the model and that region’s qMRI metrics. The correlation between the predicted and actual cell count or GO pathway scores was assessed and significant correlations were assessed at FDR-corrected p<0.05.

### Projection of PD SRT scores from qMRI data

We built multivariate linear models across all 12 control ROIs like so: cell count or GO pathway score ∼ QSM + R_1_ + R_2_^*^ `+ MTsat + PrD`. The coefficients from these models were used in combination with PD qMRI data to project PD cell counts and pathway score.

### Cook’s distance analysis of correlation robustness

For both qMRI vs SRT and predicted vs actual LOOCV correlations, Cook’s distance was calculated to assess the impact of individual data points on the overall correlation. We defined a threshold of 4/(N-k-1), where k is the number of explanatory variables and N is the number of observations (ROIs), above which individual points were considered to have a significant impact.

### Parkinson’s disease and control qMRI differences

Median qMRI metrics were extracted from the entire DLPFC at each cortical depth as well as SWM, and ANOVAs with the form `qMRI metric ∼ age + sex + SpD*diagnosis` were fitted for each metric. Significant differences between PD and controls at a particular depth were assessed using the estimated marginal means (*emmeans, v 1.10.4*) package in R (*95*) and presented at FDR-corrected *P*<0.05. Differences between PD and controls across the whole (non-partitioned) DLPFC were examined using ANOVAs of the form `qMRI metric ∼ age + sex + diagnosis`, and presented at FDR-corrected P<0.05.

### Statistical comparison of Parkinson’s disease and control linear models

We compared the linear models for Parkinson’s disease and controls qMRI predictions relative to SRT scores and counts. First, we calculated a Z-test score to have a quantitative metric of similarity of the linear fit:

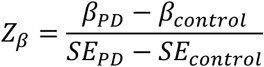

where β is the slope coefficient and SE is the standard error of the linear fit. Similarly, for the intercept:

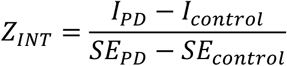

We then performed ordinary least squares regression using the python statsmodels OLS implementation. This allowed us to assess whether the differences in the β and intercept of the linear fit for controls and PD were statistically attributable to diagnosis, or whether the variation was likely to be caused by random variation, where P<0.05 were considered nominally significant and FDR-corrected p-values<0.05 were considered significant.

### GSL and GlcCer quantification

#### Glycosphingolipid (GSL) and Glucosylceramide (GlcCer) analysis

Frozen brain samples were weighed and homogenised in ddH_2_O. Protein concentration of homogenates was assayed using Pierce BCA Protein Assay (Life Technologies), and samples were prepared for extraction containing equal amounts of protein. Glycosphingolipids (GSLs) from brain homogenates were extracted and purified with chloroform: methanol (1:2, v/v) and C18 chromatography columns (Telos, Kinesis, UK). GSLs were dried down under nitrogen. Two-thirds were digested with recombinant Endoglycoceramidase I (rEGCaseI, GenScript) to obtain oligosaccharides, while one-third was digested with Cerezyme® (Genzyme, Cambridge, MA), obtaining glucose from glucosylceramide (GlcCer). The liberated free glycans and glucose were then fluorescently labelled with anthranillic acid (2AA). To remove excess 2AA label, labelled glycans and glucose were purified using DPA-6S SPE columns (Supelco, PA, USA). Purified 2AA-labelled oligosaccharides and glucose were separated and quantified by normal-phase high-performance liquid chromatography (NP-HPLC) (*97*). The NP-HPLC system consisted of a Waters Alliance 2695 separations module and an in-line Waters 2475 multi λ-fluorescence detector set at Ex λ360 nm and Em λ425 nm. The solid phase used was a 4.6 × 250 mm TSK gel-Amide 80 column (Anachem, Luton, UK). A 2AA-labelled glucose homopolymer ladder (Ludger, UK) was included to determine the glucose unit (GUs) values for the HPLC peaks. Individual GSL species were identified by their GU values and quantified by comparison of integrated peak areas with a known amount of 2AA-labelled BioQuant chitotriose standard (Ludger, UK). Results were normalised to total protein content. Due differences in processing and units, analyses for GSL and GlcCer were performed separately using a paired t-test, after which results were FDR-corrected per disease group where appropriate.

### Single nucleus RNA sequencing

#### Oligodendrocyte counts with single nuclear RNA sequencing

Public human postmortem single nuclear RNA sequencing (snRNA-seq) and single nucleus ATAC-seq data was accessed from a Parkinson’s disease study characterising early-mid stage disease (Braak 3/4), although only the snRNA-seq component was used for this study. This dataset included three cortical regions (frontal, parietal and cingulate cortex). We used preprocessing steps consistent with the original publication of the dataset (*48*). In brief, genome alignment was performed on sequencing outputs using cellranger-ARC (10x Genomics, v2.0.1) using reference genome ENSEMBL (v107, generated with cellranger-ARC v2.0.1) (*98*). Outputs were then aggregated using cellranger-ARC function `aggr`. The aggregated data was then ingested to panpipes for QC, preprocessing and clustering (*99*). Following inspection of QC metrics, nuclei were filtered based on whether they passed conventional QC metrics, namely, percentage of mitochondrial transcripts in RNA counts (<5%), percentage of ribosomal transcripts in RNA counts (<5%), percentage of haemoglobin genes (<5%), total UMI counts (<20,000), number of genes by counts (20<n<17500), number of cells per gene (>20). The data was then normalised, followed by log1p transformation. Dimensionality reduction was applied (N_PCs_ = 100) and batch correction was applied on gene-expression. Finally, community detection with the Leiden algorithm was performed.

Major cell types were detectable with cell type markers (Astrocytes: *AQP4*, *GFAP*; EndoMural cells: *CLDN5*; Neurons: *GABRB2*; Inhibitory neurons: *GAD2*; Immune cells: *CD74*, *PTPRC*, *TREM2*, *APOE*; Oligodendrocytes: *PLP1*, *MBP*; OPCs: *PDGFRA*, *BCAS1*). Each cell type was batch corrected separately using the same batch correction and community detection approach. Using cluster hierarchy, leiden resolutions were chosen per cell type based on clustering stability. Following all QC and annotation steps, a total of 740,157 nuclei remained. snRNA-seq data were then pseudobulked for downstream analyses, since this addresses pseudoreplication bias in single nuclear data (*100*). The `aggregateToPseudoBulk` function from the *dreamlet* R package v0.99.16 (*101*) was used to sum gene expression counts of nuclei by participant ID, brain region (cingulate, frontal, parietal) and cell type annotation. All covariate correction for subsequent analysis was performed consistently with the original publication.

#### Single nucleus RNA sequencing cell type proportion analysis with crumblr

For the cross-group analysis, differences in cell type proportions across groups were assessed using *crumblr* (v 0.99.6) (*102*). Briefly, occurrences of each cell type in each sample were summed. The data was covariate corrected (excluding binary variables, making the final variables: sex, batch, total_deduplicated_percentage) using `dream()` from *variancePartition* (v 1.32.5) (*103*) and the fit was smoothed using eBayes() from *limma* (v 3.58.1) (*104*). This comparison was performed separately for each brain region analysed, and pairwise comparisons were performed for controls vs PD. Finally, p-values were FDR-corrected.

### Bulk RNA sequencing

#### Post-mortem sample selection

All brain samples used in this study were provided by the Parkinson’s Disease UK tissue bank with informed consent to use the material for research available for each donation. Selection of cases and controls was performed based upon disease status of each individual (Control or Parkinson’s disease) found in the clinical notes in patient records, no dementia incidence, post-mortem delay (less than 24 hours) and an absence of any confounding pathology. The cases with the highest RIN were selected from those that met these criteria. Formalin-fixed paraffin-embedded sections and frozen sections or chips of tissue were taken as necessary for all further downstream analyses and stored at room temperature or –80°C respectively until use.

Only those that were reported not to have dementia and mapped to the Brainstem predominant or limbic transitional stage were selected. PD cases had no confounding pathology, but some controls did have some minimal signs of age-related changes. The two cohorts, namely control and PD, had comparable mean age of death (agecontrols = 80.4; agePD = 79.5) and both groups had a higher representation of male subjects relative to female (ratiocontrols = 1.857; ratioPD = 2.0).

#### Bulk RNA isolation

Approximately 50-100 g of frozen tissue was used per brain sample for RNA extraction. Tissue lysis was performed using QIAzol and RNA was extracted using the RNeasy 96 Kit (Qiagen) with an on-membrane DNase treatment, as per manufacturer instructions. Samples were quantified by absorption on the QIAxpert (Qiagen), and RNA integrity number (RIN) measured using the Agilent 4200 Tapestation (Agilent Technologies).

#### Generation of bulk RNA sequencing data

Stranded cDNA libraries were constructed with the KAPA mRNA HyperPrep Kit (Roche) used as per manufacturer instructions. More specifically, 500ng of total RNA was used as input for each sample and xGen Dual Index UMI Adapters (Integrated DNA Technologies, Inc.) were added to each sample to minimise read mis-assignment when performing post-sequencing sample de-multiplexing. Libraries were multiplexed on S2 and S4 NovaSeq flow cells for paired-end 150 bp sequencing on the NovaSeq 6000 Sequencing System (Illumina) to obtain a target read depth of 110 million paired-end reads per sample. Sequenced reads were de-multiplexed and FASTQ files were generated using the BCL Convert software (Illumina). FASTQ files were run through a *Nextflow* pipeline (*105*) available at https://github.com/Jbrenton191/RNAseq_splicing_pipeline. Briefly, this *Nextflow* pipeline performed quality control (QC) of reads and aligned the samples to the genome and transcriptome (GRCh38 and Gencode v41 annotations). Briefly, initial QC was performed using *fastp* (v 0.23.2) (*106*), which removed low quality reads and bases, as well as the trimming of adapter sequences. Filtered reads were aligned to the transcriptome using *Salmon* v1.9.0 (*107*) correcting for sequence-specific, fragment GC-content and positional biases; using the entire genome as a decoy sequence. For splicing analyses, reads were aligned to the genome using *STAR* (v 2.7.10a, with two-pass mode) (*108*). *FastQC* (v0.11.9) (*109*) was used to generate read QC data before and after *fastp* (*106*) filtering. *RSEQC* (v4.0.0) (*110*), *Qualimap* (v2.2.2a) (*111*) and *Picard* (v2.27.5) (https://github.com/broadinstitute/picard) were used to generate alignment QC metrics. *MultiQC* (v1.13) (*112*), was used to visualise and collate quality metrics from all pipeline modules.

#### Sample quality control, covariate selection and outlier removal

Samples with a low read count (<60M reads aligning to the transcriptome), a high % of reads aligning to intronic regions of the genome (>30%) or a deviant average GC content (<40% or >65%) were removed from further analysis. Transcriptome aligned reads generated from *Salmon* were used to identify covariates and then outlier samples. For both processes, salmon read files were imported into R (v4.3.0) and converted into gene-level count matrices using the *tximport* (v1.30.0) (*113*) and *DESeq2* (v1.42.1) (*114*) packages.

Covariates and outliers were detected with the same method as above. The covariate selection was run once across all bulk samples and covariates passing the selection criteria for all samples were taken forwards. The outlier analysis was run using these covariates and z-scores were calculated for each sample against the sample population from the same brain area.

#### Differential gene expression analysis

Gene-level count matrices were filtered for genes with 0 counts in at least one sample in either the disease or control group. Analysis was performed using *limma*/*voom* (*104*, *115*). Briefly, *voom* was used to shrink the dispersion and the `duplicateCorrelation()` function was used to remove the effect of each individual to allow multiple brain areas from the same individual to be modelled together. These functions were applied twice as recommended by the developers (https://search.r-project.org/CRAN/refmans/DGEobj.utils/html/runVoom.html). Group and brain area factors were merged into a single term to allow the model to be run once and all comparisons to be extracted from that model. Contrasts were run between groups for each brain area, and multi-region comparisons incorporating either all brain regions, cortical regions (anterior cingulate cortex, frontal cortex, parahippocampal gyrus, parietal cortex and temporal cortex) or subcortical regions (substantia nigra, putamen and caudate). Differentially expressed genes were extracted for each contrast using the `eBayes()` and `topTable()` *limma* (*104*) functions and were classified as differentially expressed if their FDR-adjusted p value was ≤ 0.05.

#### Gene set enrichment analysis

Gene set enrichment analysis was performed using the *fgsea* package (v 1.28.0) (*116*), using the gene lists from the literature, including genes expressing myelin structural proteins (*117*) and genes associated with oligodendrocyte maturation (*118*). Log2 fold changes were used to rank all genes, both significant and non-significant, and the `fgsea()` function was applied for the combined brain area comparison with `nPermSimple = 5000`. P-values were corrected by the number of gene lists tested (n=2).

**Supplementary Figure 1.**
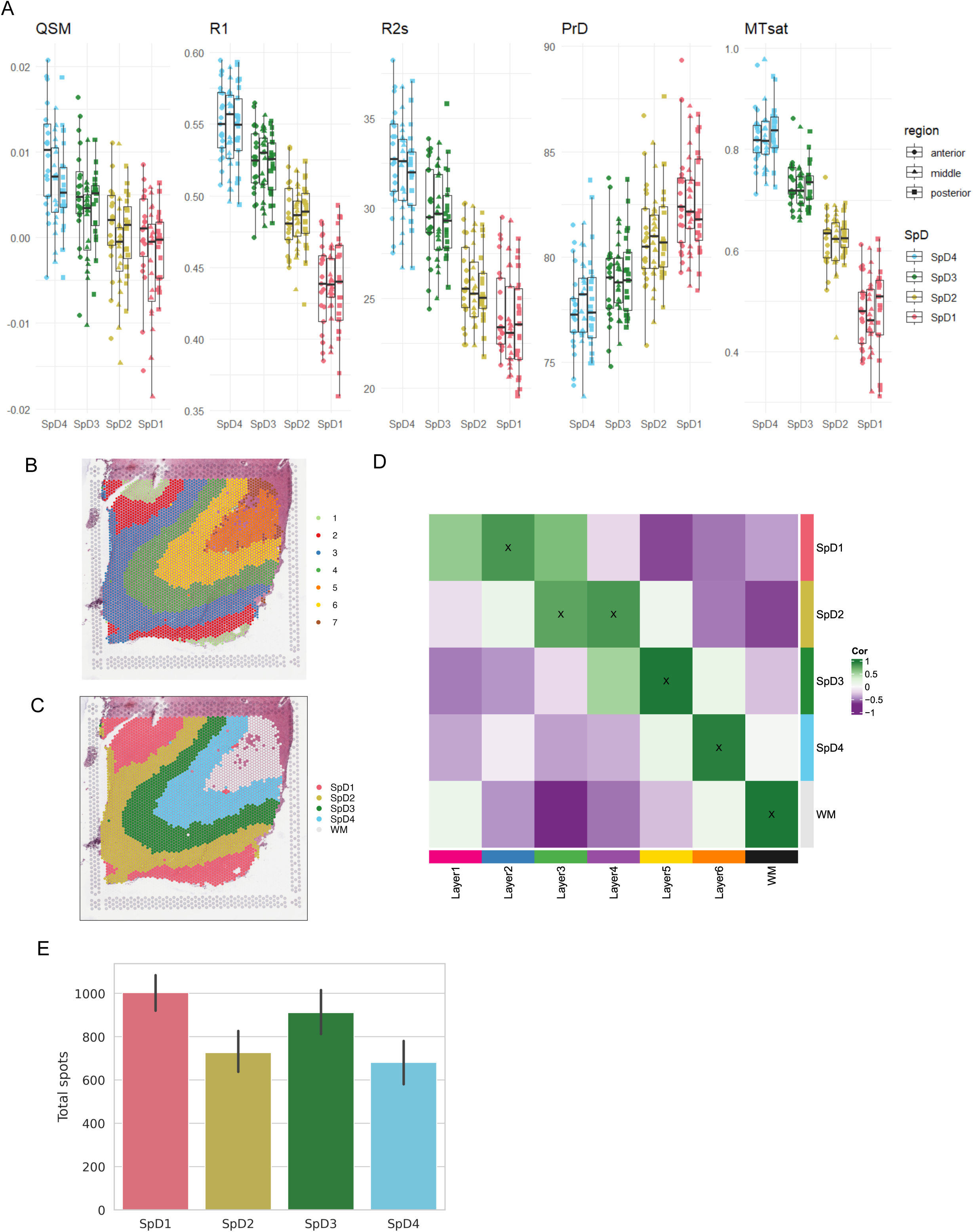
– Spatially resolved transcriptomic data in DLPFC. (**A**) Median qMRI metric signals for all ROIs in control DLPFC, adjusting for age and sex. (**B**) Spot plot of representative section Br2743_mid showing (i) k=7 clustering with *BayesSpace* in relation to MRI compatible annotation of (ii) four cortex SpDs and white matter (WM). (**C**) *spatialLIBD* (*91*) spatial registration heatmap of SRT DLPFC data (*34*) with four SpDs and WM compared to reference SRT data of DLPFC with layer annotations (*92*), summarizing correlations between enrichment *t*-statistics from each dataset. High correlation (green) represents similar patterns of gene expression, highest confidence matches are highlighted with an “X”. (**D**) Bar plot of total number of spots per cortex SpD in SRT data.

**Supplementary Figure 2.**
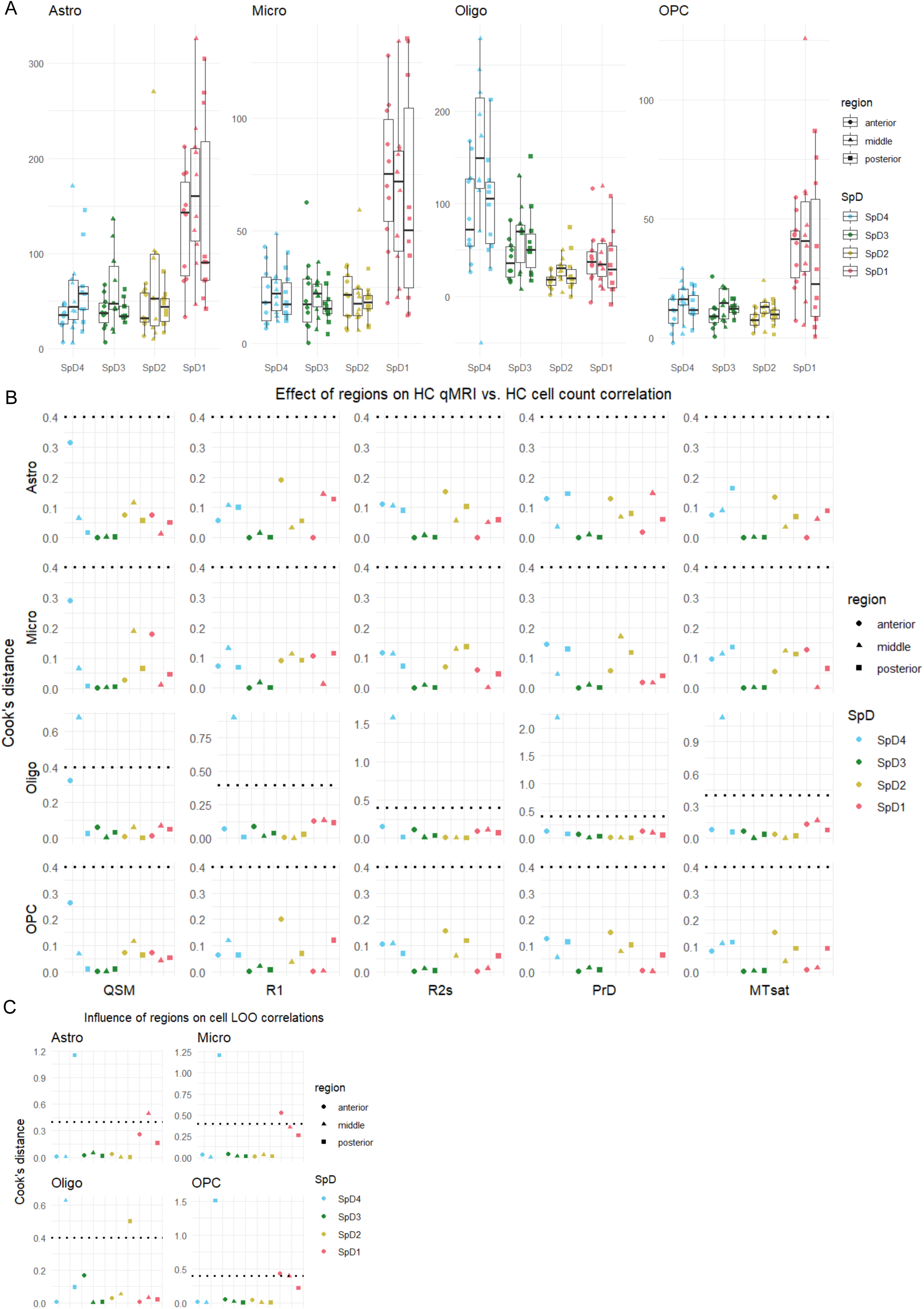
– SRT deconvoluted control cell counts and predicted count robustness. (**A**) SRT-deconvoluted cell counts (y-axis) per ROI (x-axis) in control DLPFC. (**B**) Cook’s distance plots showing robustness of correlations between cell type counts and qMRI metrics.The dotted lines indicate a threshold of 4/(N-k-1), where k is the number of explanatory variables and N is the number of observations (ROIs), above which removal of individual points was considered to have a significant impact on the correlation. The correlations were generally robust to the removal of any one ROI. (**C**) Cook’s distance plots showing level of robustness of correlations between predicted cell counts and control SRT-deconvoluted cell counts. Astrocyte, microglia and OPC correlations were vulnerable to the removal of select SpD_1_ and SpD_4_ ROIs, while oligodendrocyte correlations were vulnerable to the removal of select SpD_2_ and SpD_4_ ROIs. SpD = spatial domain; QSM = quantitative susceptibility mapping; R_1_ = longitudinal relaxation rate; R2s = R_2_*, transverse relaxation rate; PrD = proton density; MTsat = magnetisation transfer saturation; SpD = spatial domain; QSM = quantitative susceptibility mapping; R2s = R_2_*, transverse relaxation rate; PrD = proton density; Astro = astrocytes; Micro = microglia; Oligo = oligodendrocytes; OPC = Oligodendrocyte progenitor cells.

**Supplementary Figure 3.**
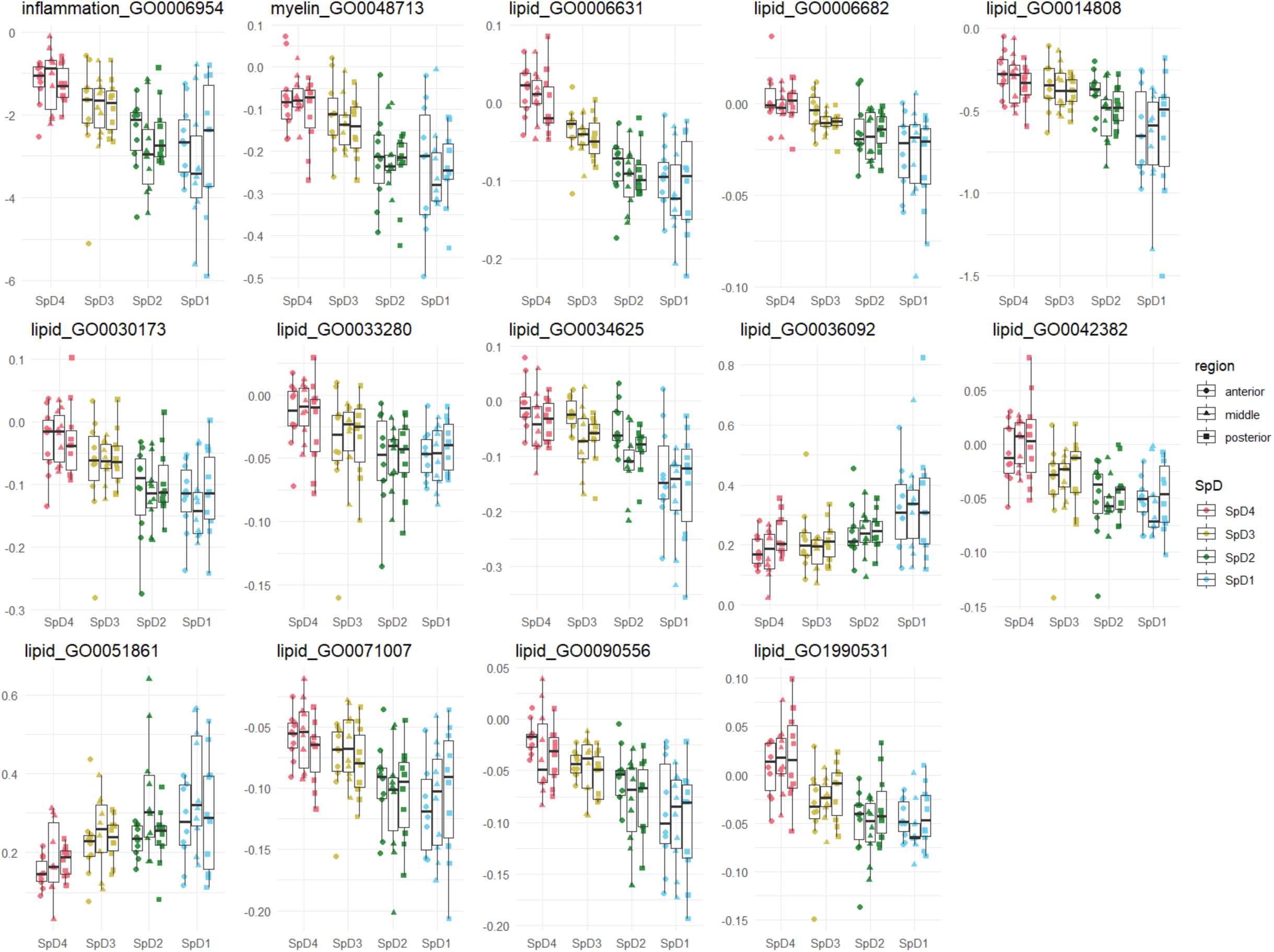
– Association between control median qMRI metrics and control median GO pathway scores across cortical spatial domains. Median GO pathway scores (y-axis) over ROIs (x-axis). Pathways that have R^2^ > 0.85 and pFDR < 0.05 with at least one qMRI metric are displayed. Both GO pathway scores and qMRI metrics are adjusted for age and sex. Anterior, middle and posterior refer to subregions of the dorsolateral prefrontal cortex. SpD = spatial domain; QSM = quantitative susceptibility mapping; R_1_ = longitudinal relaxation rate; R2s = R_2_*, transverse relaxation rate; PrD = proton density; MTsat = magnetisation transfer saturation; inflammation_GO0006954 = inflammatory response; myelin_GO0048713 = regulation of oligodendrocyte differentiation; lipid_GO0006631 = fatty acid metabolic process; lipid_GO0006682 = galactosylceramide biosynthetic process; lipid_GO0014808 = release of sequestered calcium ion into cytosol by sarcoplasmic reticulum; lipid_GO0030173 = obsolete integral component of Golgi membrane; lipid_GO0033280 = response to vitamin D; lipid_GO0034625 = fatty acid elongation, monounsaturated fatty acid; lipid_GO0036092 = phosphatidylinositol-3-phosphate biosynthetic process; lipid_GO0042382 = paraspeckles; lipid_GO0051861 = glycolipid binding; lipid_GO0071007 = U2-type catalytic step 2 spliceosome; lipid_GO0090556 = phosphatidylserine floppase activity; lipid_GO1990531=phospholipid-translocating ATPase complex.

**Supplementary Figure 4.**
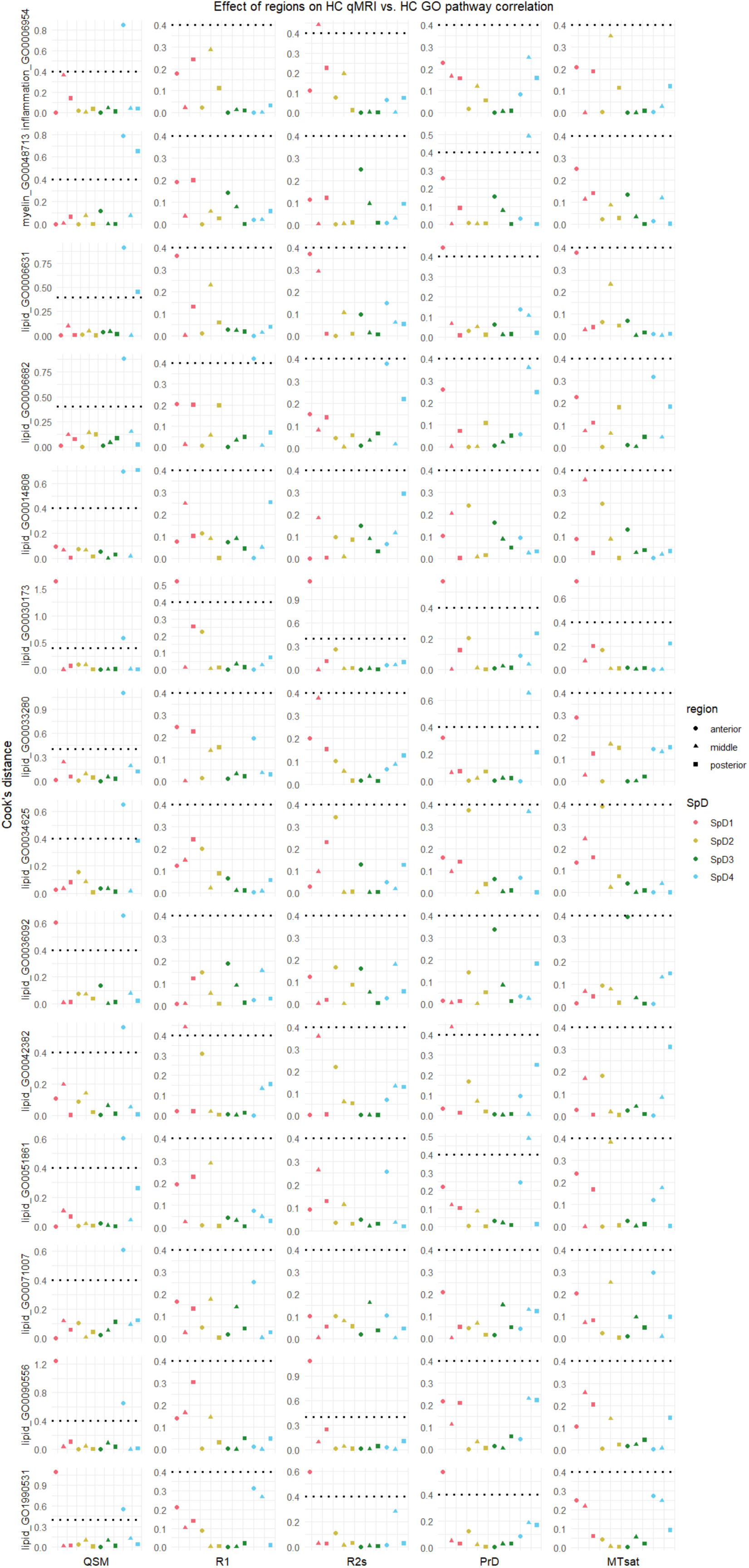
– Cook’s distance assessment of control correlation robustness for GO pathways in relation to qMRI metrics. Most correlations were robust to the removal of specific ROIs (41/70 robust to all, 22/70 to all but one, 7 to all but two). SpD = spatial domain; QSM = quantitative susceptibility mapping; R1 = longitudinal relaxation rate; R2s = R_2_*, transverse relaxation rate; PrD = proton density; MTsat = magnetisation transfer saturation; inflammation_GO0006954 = inflammatory response; myelin_GO0048713 = regulation of oligodendrocyte differentiation; lipid_GO0006631 = fatty acid metabolic process; lipid_GO0006682 = galactosylceramide biosynthetic process; lipid_GO0014808 = release of sequestered calcium ion into cytosol by sarcoplasmic reticulum; lipid_GO0030173 = obsolete integral component of Golgi membrane; lipid_GO0033280 = response to vitamin D; lipid_GO0034625 = fatty acid elongation, monounsaturated fatty acid; lipid_GO0036092 = phosphatidylinositol-3-phosphate biosynthetic process; lipid_GO0042382 = paraspeckles; lipid_GO0051861 = glycolipid binding; lipid_GO0071007 = U2-type catalytic step 2 spliceosome; lipid_GO0090556 = phosphatidylserine floppase activity; lipid_GO1990531 = phospholipid-translocating ATPase complex; SpD = spatial domain; QSM = quantitative susceptibility mapping; R2s = R_2_*, transverse relaxation rate; PrD = proton density.

**Supplementary Figure 5.**
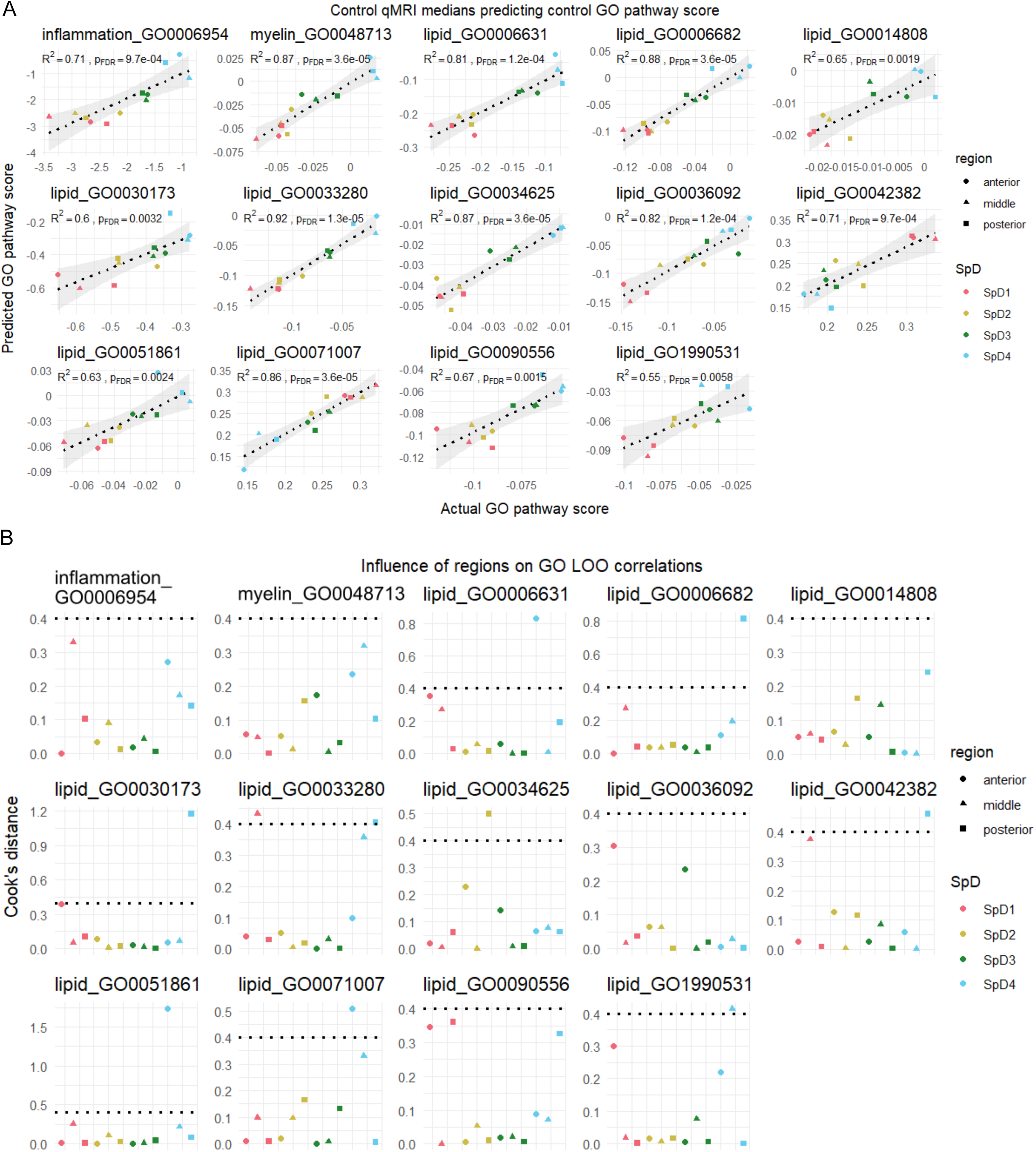
– Assessment of control GO pathway prediction score accuracy and robustness. (**A**) Correlative analysis of predicted pathway scores relative to SRT-based pathway scores, all correlations are statistically significant. (**B**) Cook’s distance statistical analysis to test robustness of correlations when removing each ROI. Inflammation and myelin pathways were robust to the removal of all ROIs, whilst lipid pathways were sensitive to the removal of some ROIs. SpD = spatial domain; QSM = quantitative susceptibility mapping; R_1_ = longitudinal relaxation rate; R2s = R_2_*, transverse relaxation rate; PrD = proton density; MTsat = magnetisation transfer saturation; inflammation_GO0006954 = inflammatory response; myelin_GO0048713 = regulation of oligodendrocyte differentiation; lipid_GO0006631 = fatty acid metabolic process; lipid_GO0006682 = galactosylceramide biosynthetic process; lipid_GO0014808 = release of sequestered calcium ion into cytosol by sarcoplasmic reticulum; lipid_GO0030173 = obsolete integral component of Golgi membrane; lipid_GO0033280 = response to vitamin D; lipid_GO0034625 = fatty acid elongation, monounsaturated fatty acid; lipid_GO0036092 = phosphatidylinositol-3-phosphate biosynthetic process; lipid_GO0042382 = paraspeckles; lipid_GO0051861 = glycolipid binding; lipid_GO0071007 = U2-type catalytic step 2 spliceosome; lipid_GO0090556 = phosphatidylserine floppase activity; lipid_GO1990531 = phospholipid-translocating ATPase complex; SpD = spatial domain; QSM = quantitative susceptibility mapping; R2s = R_2_*, transverse relaxation rate; PrD = proton density.

**Supplementary Figure 6.**
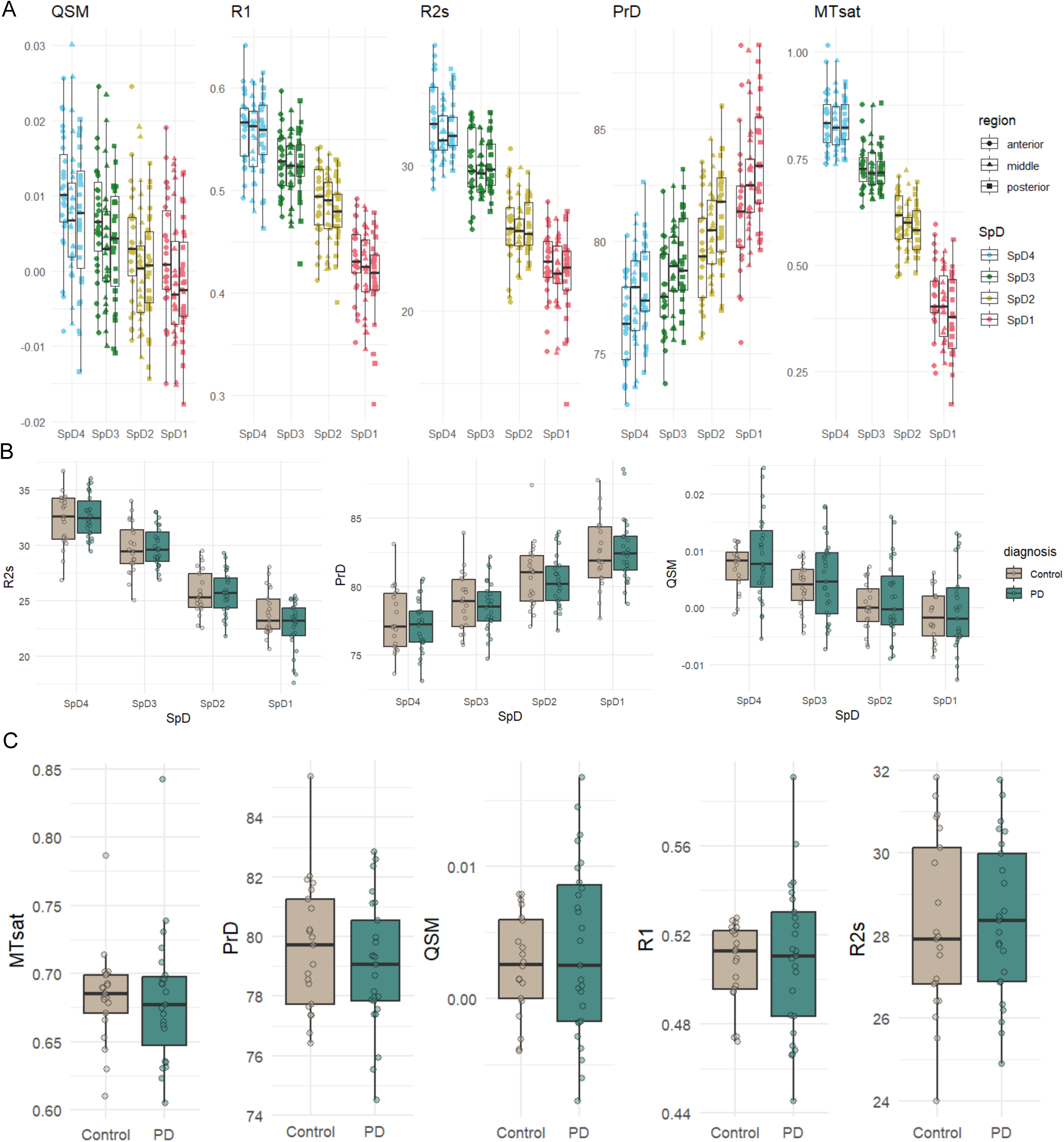
– qMRI spatial characterisation of Parkinson’s disease. (**A**) Boxplots showing median qMRI signals across all ROIs in PD cases. Values are adjusted for participant age and sex. (**B**) Comparison of qMRI metrics between PD and controls for R_2_*, PrD, and QSM. ANOVAs adjusting for age and sex revealed no significant differences between control and PD in any SpD at pFDR < 0.05. SpD = spatial domain. (**C**) Comparison of qMRI metrics between PD and controls across the DLPFC when partitioned into spatial domains. ANOVAs adjusting for age and sex revealed no differences at pFDR < 0.05 or at p<0.05. QSM = quantitative susceptibility mapping; R_1_ = longitudinal relaxation rate; R2s = R_2_*, transverse relaxation rate; PrD = proton density; MTsat = magnetisation transfer saturation; SpD = spatial domain; QSM = quantitative susceptibility mapping; R2s = R_2_*, transverse relaxation rate; PrD = proton density.

**Supplementary Figure 7.**
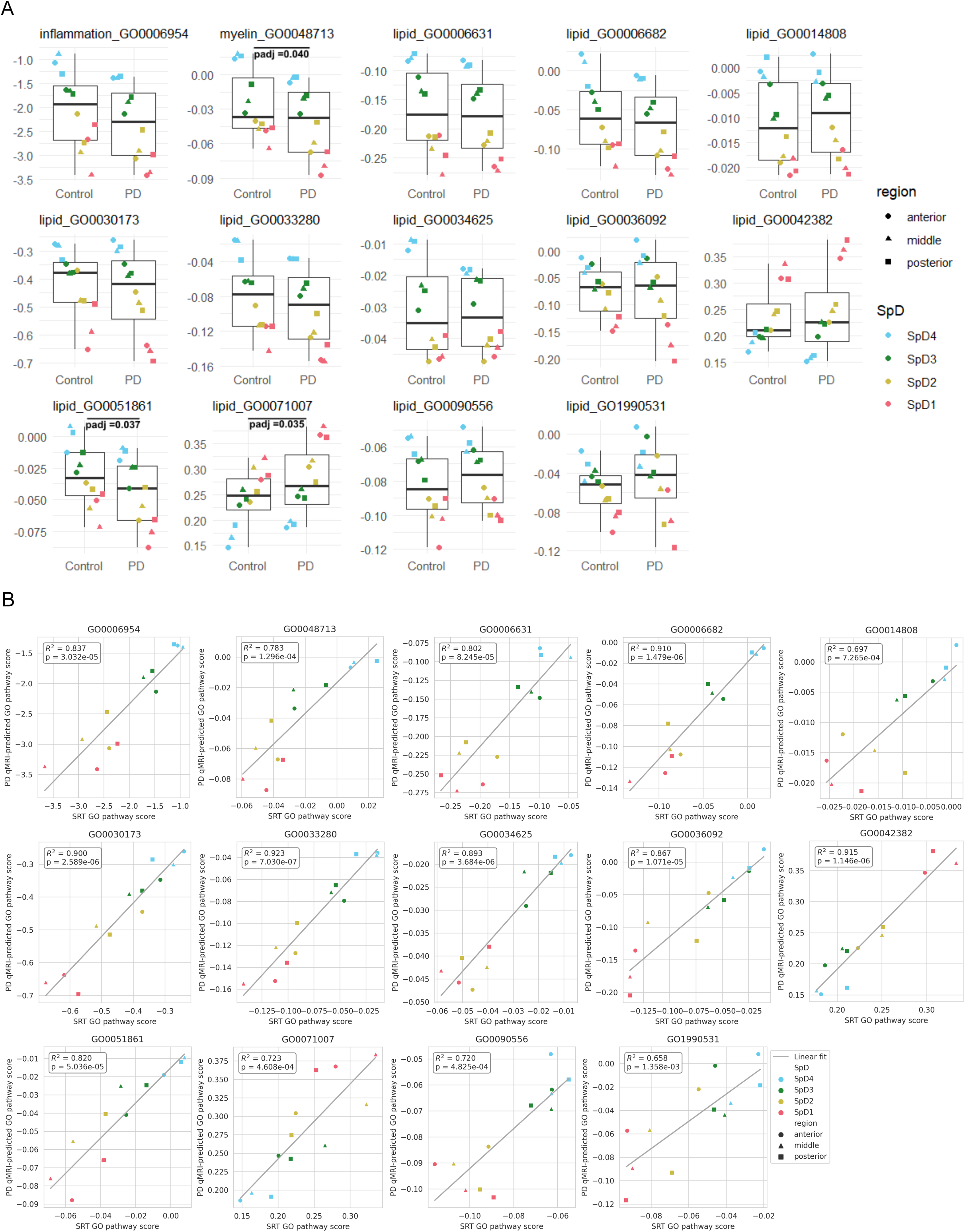
– Prediction of control and PD molecular pathway information from qMRI data. (**A**) Comparison of overall predicted pathway scores in PD to control (HC) pathway scores calculated with SRT. None of these pathways were predicted to have significant differences in PD. (**B**) Correlation of predicted pathway scores in PD relative to real pathway scores in control SRT. All correlations are statistically significant, suggesting there are no expected spatial shifts in the expression of these pathways. SpD = spatial domain; QSM = quantitative susceptibility mapping; R_1_ = longitudinal relaxation rate; R2s = R_2_*, transverse relaxation rate; PrD = proton density; MTsat = magnetisation transfer saturation; inflammation_GO0006954 = inflammatory response; myelin_GO0048713 = regulation of oligodendrocyte differentiation; lipid_GO0006631 = fatty acid metabolic process; lipid_GO0006682 = galactosylceramide biosynthetic process; lipid_GO0014808 = release of sequestered calcium ion into cytosol by sarcoplasmic reticulum; lipid_GO0030173 = obsolete integral component of Golgi membrane; lipid_GO0033280 = response to vitamin D; lipid_GO0034625 = fatty acid elongation, monounsaturated fatty acid; lipid_GO0036092 = phosphatidylinositol-3-phosphate biosynthetic process; lipid_GO0042382 = paraspeckles; lipid_GO0051861 = glycolipid binding; lipid_GO0071007 = U2-type catalytic step 2 spliceosome; lipid_GO0090556 = phosphatidylserine floppase activity; lipid_GO1990531 = phospholipid-translocating ATPase complex; SpD = spatial domain.

**Supplementary Figure 8.**
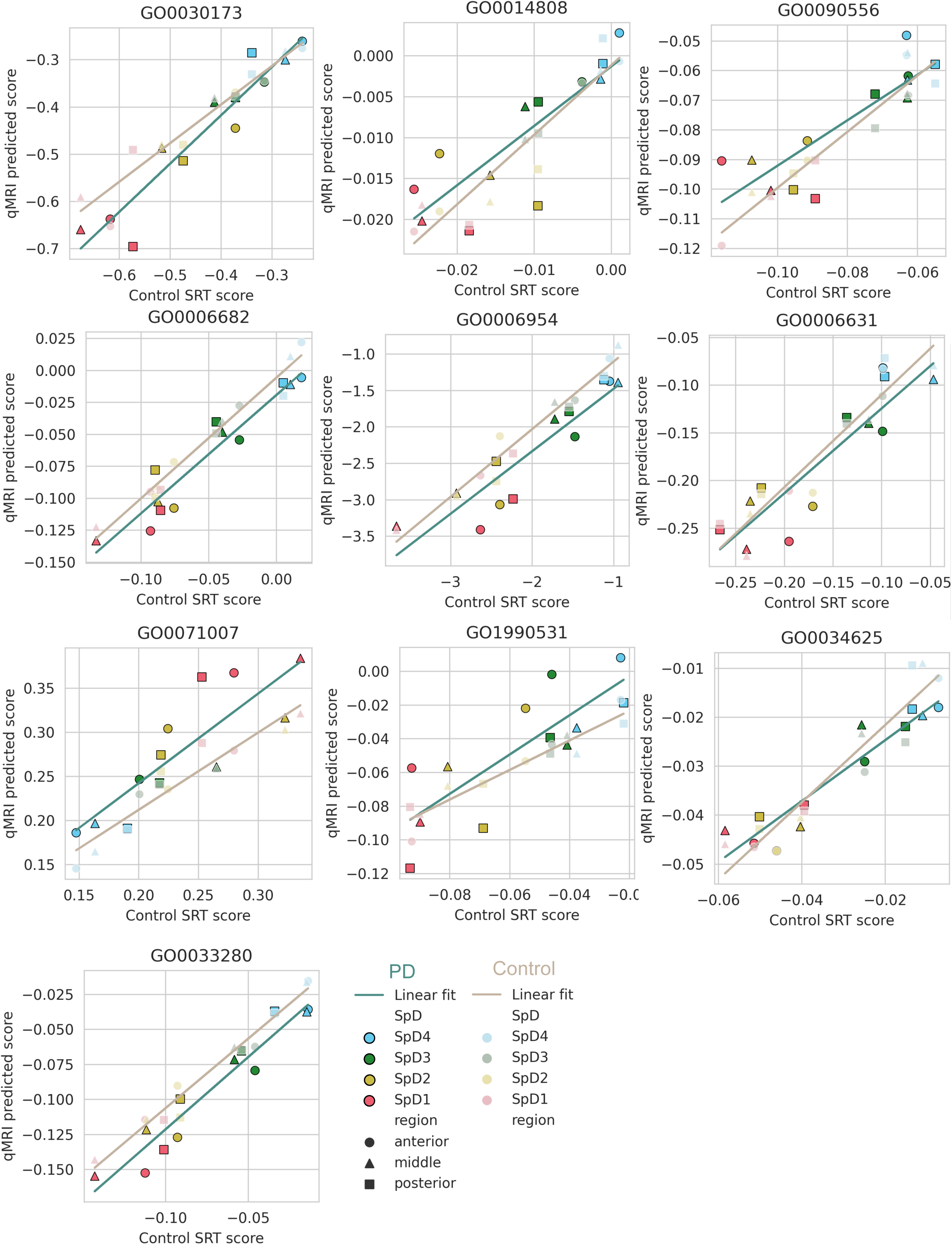
– Overlaid correlation of SRT score vs predicted pathway score for controls and PD. Pathways where diagnosis does not impact the model (8.0×10^−2^ < p-value < 9.6×10^−2^). SpD = spatial domain; QSM = quantitative susceptibility mapping; R_1_ = longitudinal relaxation rate; R2s = R_2_*, transverse relaxation rate; PrD = proton density; MTsat = magnetisation transfer saturation; inflammation_GO0006954 = inflammatory response; lipid_GO0006631 = fatty acid metabolic process; lipid_GO0006682 = galactosylceramide biosynthetic process; lipid_GO0014808 = release of sequestered calcium ion into cytosol by sarcoplasmic reticulum; lipid_GO0030173 = obsolete integral component of Golgi membrane; lipid_GO0033280 = response to vitamin D; lipid_GO0034625 = fatty acid elongation, monounsaturated fatty acid; lipid_GO0071007 = U2-type catalytic step 2 spliceosome; lipid_GO0090556 = phosphatidylserine floppase activity; lipid_GO1990531 = phospholipid-translocating ATPase complex; SpD = spatial domain.

**Supplementary Figure 9.**
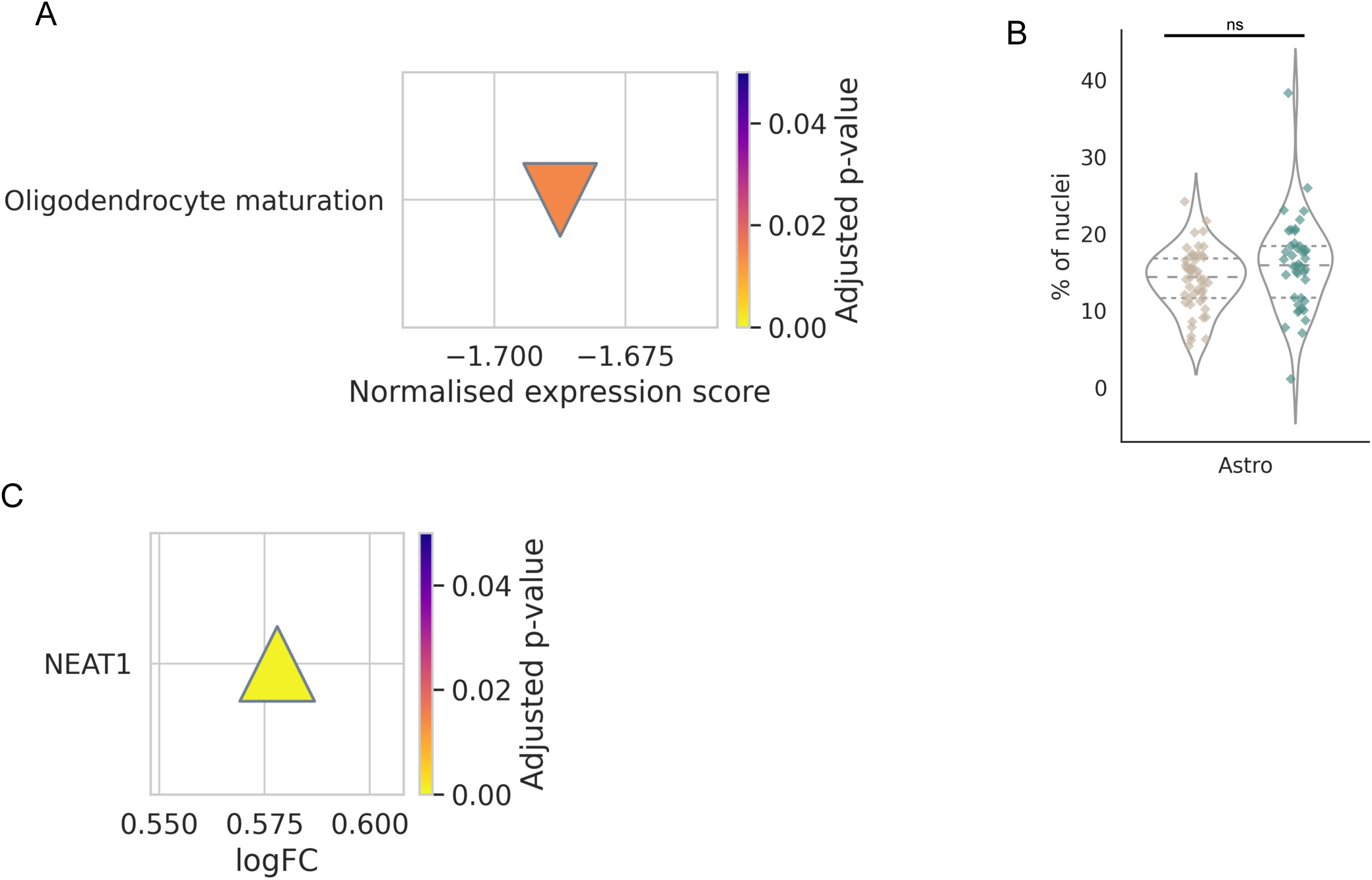
– Transcriptomic data supports changes to predicted pathways. (A) In bulk RNA-seq, gene set enrichment analysis shows that genes associated with oligodendrocyte maturation are significantly less expressed in PD than in controls across all regions in bulk RNA-seq dataset (enrichment score = –1.68, FDR-adjusted p-value = 2.45×10^−2^). (B) Cell type proportion analysis comparing the proportion of astrocytes in PD and controls show there are no significant changes across the overall population of astrocytes (**C**) Differential gene expression of *NEAT1*, an essential structural scaffold for paraspeckle formation, in bulk RNA-seq of anterior cingulate cortex (logFC = 0.578, FDR-adjusted p-value = 7.9×10^−4^).

**Supplementary Figure 10.**
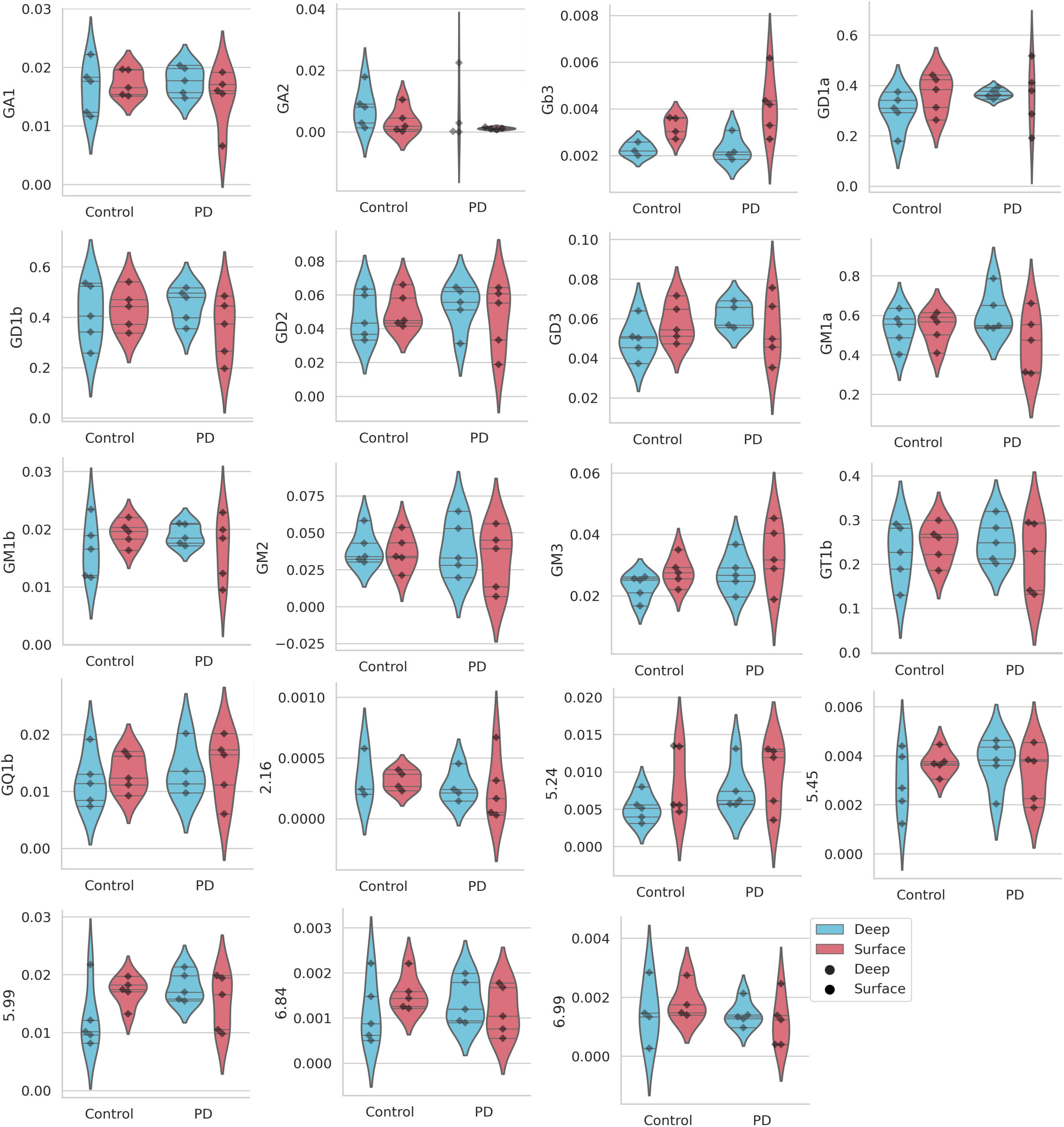
– Glycosphingolipid quantification in deep and surface cortex from control and PD postmortem brain samples. Intra-individual pair-wise comparisons of deep and surface GSL levels did not find any statistically significant differences in concentration following FDR correction. Deep and surface cortex differences in Gb3 were nominally significant for PD (p-value = 1.9 x 10^−2^).

